# Rewinding the ratchet: rare recombination locally rescues neo-W degeneration and generates plateaus of sex-chromosome divergence

**DOI:** 10.1101/2024.01.20.576444

**Authors:** Thomas Decroly, Roger Vila, Konrad Lohse, Alexander Mackintosh

## Abstract

Natural selection is less efficient in the absence of recombination. As a result, non-recombining sequences, such as sex chromosomes, tend to degenerate over time. Although the outcomes of recombination arrest are typically observed after many millions of generations, recent neo-sex chromosomes can give insight into the early stages of this process. Here we investigate the evolution of neo-sex chromosomes in the Spanish marbled white butterfly, *Melanargia ines*, where a Z-autosome fusion has turned the homologous autosome into a non-recombining neo-W chromosome. We show that these neo-sex chromosomes are likely limited to the Iberian population of *M. ines*, and that they arose around the time when populations in Iberia and North-Africa split, around 1.5 million years ago. Recombination arrest of the neo-W chromosome has led to an excess of premature stop codons and frameshift mutations, while levels of gene expression have remained similar for the neo-W and neo-Z chromosomes, even for genes with loss-of-function mutations. Surprisingly, we identified two regions of *∼* 1 Mb at one end of the neo-W that are both less diverged from the neo-Z and less degraded than the rest of the chromosome, suggesting a history of rare but repeated genetic exchange between the two neo-sex chromosomes. These plateaus of neo-sex chromosome divergence suggest that neo-W degradation can be locally reversed by rare recombination between neo-W and neo-Z chromosomes.

## Introduction

Recombination allows alleles to move between genetic backgrounds. This results in more efficient natural selection, as the fitness effect of a new allele is decoupled from the background on which it arose (Fisher 1930). A reduction in the rate of recombination therefore results in less efficient purging of deleterious and reduced fixation of beneficial alleles (Muller 1964, Hill & Robertson 1968). While recombination is often considerably reduced in particular regions of the genome, e.g. nearby centromeres (Choo 1998), it can also be suppressed entirely on certain chromosomes. Sex-limited chromosomes may only recombine at a particular region (e.g. the pseudoautosomal region of the mammalian Y chromosome) or not at all (e.g. W chromosomes of Lepidoptera). Non-recombining sex chromosomes typically exhibit signs of sequence degeneration such as elevated rates of loss-of-function mutations, sequence loss, TE proliferation and pseudogenisation of genes (Bachtrog 2013). Genomic analyses of non-recombining sex chromosomes show that recombination arrest leads to natural selection becoming less efficient, but it is less clear exactly which evolutionary processes (e.g. Muller’s ratchet, hitchhiking of deleterious mutations, background selection) are most important in this process (Charlesworth & Charlesworth 2000, Bachtrog 2008). There are also unresolved questions about how non-recombining chromosomes affect gene expression, and, ultimately, phenotypes. For example, it is unclear whether the low levels of gene expression observed on some non-recombining chromosomes come about gradually as a result of sequence degeneration, or if instead functional genes are silenced first, allowing for the neutral accumulation of loss-of-function mutations (Lenormand et al. 2020). Resolving these questions requires both careful inference from natural examples of recombination arrest and population genetic modelling of the relevant processes (Bachtrog 2008).

Neo-sex chromosomes that form as a result of a fusion between an autosome and a sex chromosome have been reported in a wide variety of taxa (Howell et al. 2009, Gil-Ferńandez et al. 2020, Huang et al. 2022, Akagi et al. 2023, Sacchi et al. 2023). Sex-autosome linkage means that the previously autosomal chromosome will follow a sex-specific pattern of inheritance. Consequently, in species with achiasmatic meiosis (where recombination only happens in the homogametic sex) the evolution of neo-sex chromosomes leads to recombination suppression. In *Drosophila*, where meiosis in males is typically achiasmatic, a neo-Y chromosome will experience a sudden arrest of recombination. The same is also true for neo-W chromosomes in Lepidoptera, as female meiosis is achiasmatic. Recombination arrest can also be generated by X-autosome and Z-autosome fusions, as the unfused autosome will co-segregates with the Y / W chromosome, effectively forming non-recombining neo-Y / neo-W chromosomes. Recent sex-autosome chromosome fusions in achiasmate species can thus provide insight into the effect of recombination suppression (Charlesworth & Charlesworth 2000, Wei & Bachtrog 2019).

In Lepidoptera, the Z chromosome is involved in fusions more often than any autosome (Wright et al. 2023), leading to a high rate of sex chromosome evolution. While many of the neo-sex chromosomes known in Lepidoptera are old (i.e. shared by multiple genera), young neo-sex chromosomes have been described in a handful of taxa (Smith et al. 2016, Mongue et al. 2017, Mackintosh et al. 2022, Rueda-M et al. 2023, Hööok et al. 2023). Some of these neo-sex chromosomes are nonetheless highly diverged, meaning that little can be inferred about the early stages of neo-W degeneration (Mongue et al. 2017). Others, however, are so young that little degeneration is observed at all (Martin et al. 2020). Recently, neo-sex chromosomes have been identified in the *sara* / *sapho* clade of *Heliconius* (Rueda-M et al. 2023) as well as in the pierid butterfly *Leptidea sinapis* (Hööok et al. 2023), with both cases involving step-wise fusions of autosomes to sex chromosomes. Importantly, the neo-sex chromosomes in these taxa are old enough to have accumulated some divergence but young enough to provide insight into the consequences of recombination arrest. Identification and analysis of natural systems such as these provides an opportunity to better understand how non-recombining sex chromosomes evolve over time.

The nymphalid butterfly *Melanargia ines* (the Spanish marbled white) is found on the Iberian Peninsula and the Maghreb. A previous analysis of mitochondrial sequence data revealed a deep split between Iberian and North-African populations, as well as a more recent split between Western and Eastern regions of the Maghreb (Dapporto et al. 2022). Additionally, the karyotype of *M. ines* has been reported as *n* = 13 in males (de Lesse 1970), which is far lower than that of most Nymphalidae butterflies and suggests a recent history of multiple chromosome fusions. Here, we report the discovery of a Z-autosome fusion in the Iberian population of *M. ines*. These neo-sex chromosomes are intermediate in age, and so, represent a promising system to characterise the mechanisms of sequence degeneration due to recombination arrest. We generate a chromosome-level genome assembly and analyse whole-genome resequencing (WGS) data using comparative and population genomic methods to address the following questions:

- What is the distribution of the neo-sex chromosomes across *M. ines* populations?
- What is the age and evolutionary history of the neo-sex chromosomes and how does it relate to the population history of *M. ines*?
- How complete is recombination arrest of the neo-W chromosome?
- How degenerated is the neo-W chromosome?

## Results

### A genome assembly of *Melanargia ines*

We generated a chromosome-level genome assembly for *M. ines* using a combination of Pacbio long-read, Illumina short-read and HiC data generated from two individuals sampled in Portugal (Table S1). The assembly is 421.7 Mb in length, with a contig N50 of 4.3 Mb. The assembly contains 14 chromosome-level sequences (hereafter simply referred to as chromosomes) which range from 13.0 to 51.4 Mb in length. In contrast, de Lesse (1970) only observed 13 chromosomes in the spermatocytes of a male *M. ines* individual from Teruel, Spain. We further investigated the karyotype of *M. ines* by constructing haplotype-specific HiC maps as described in Mackintosh et al. (2022). These HiC maps, which were generated from a female sample from Portugal, contain very few read pairs that map exclusively to chromosome 13. This is consistent with chromosome 13 being the Z chromosome, as this would be present in a single copy in females and so cannot be phased into haplotypes (we also confirm this result using sex-specific read coverage). We find that chromosomes 13 and 14 display a high density of HiC contacts in one haplotype, but not the other (Figure 1A, 1B). These results are consistent with this individual possessing a single Z-autosome fusion (i.e. a neo-Z) as well as another (unfused) copy of the same autosome. The Z-autosome fusion is also supported by a genome alignment between *M. ines* and a closely related species *M. galathea* (Figure S1). Given that the W chromosome is absent from our genome assembly (contigs were assembled from male-derived reads), we cannot test whether the neo-W copy of chromosome 14 is fused to the W. However, we expect this chromosome to behave as a neo-W irrespective of physical linkage to the ancestral W given that meiosis is achiasmatic in females. Assuming that the neo-Z fusion is at high frequency, we would expect males to have 2*n* = 26 chromosomes (12 pairs of autosomes + one pair of neo-Z chromosomes), whereas females should have 2*n* = 27 chromosomes (12 pairs of autosomes + one copy each of the neo-Z, the ancestral W and the neo-W). Our data is therefore consistent with the observations of de Lesse (1970) and is suggestive of a young neo-sex chromosome system given that neo-Z and neo-W reads both map to the neo-Z reference sequence.

**Figure 1.**
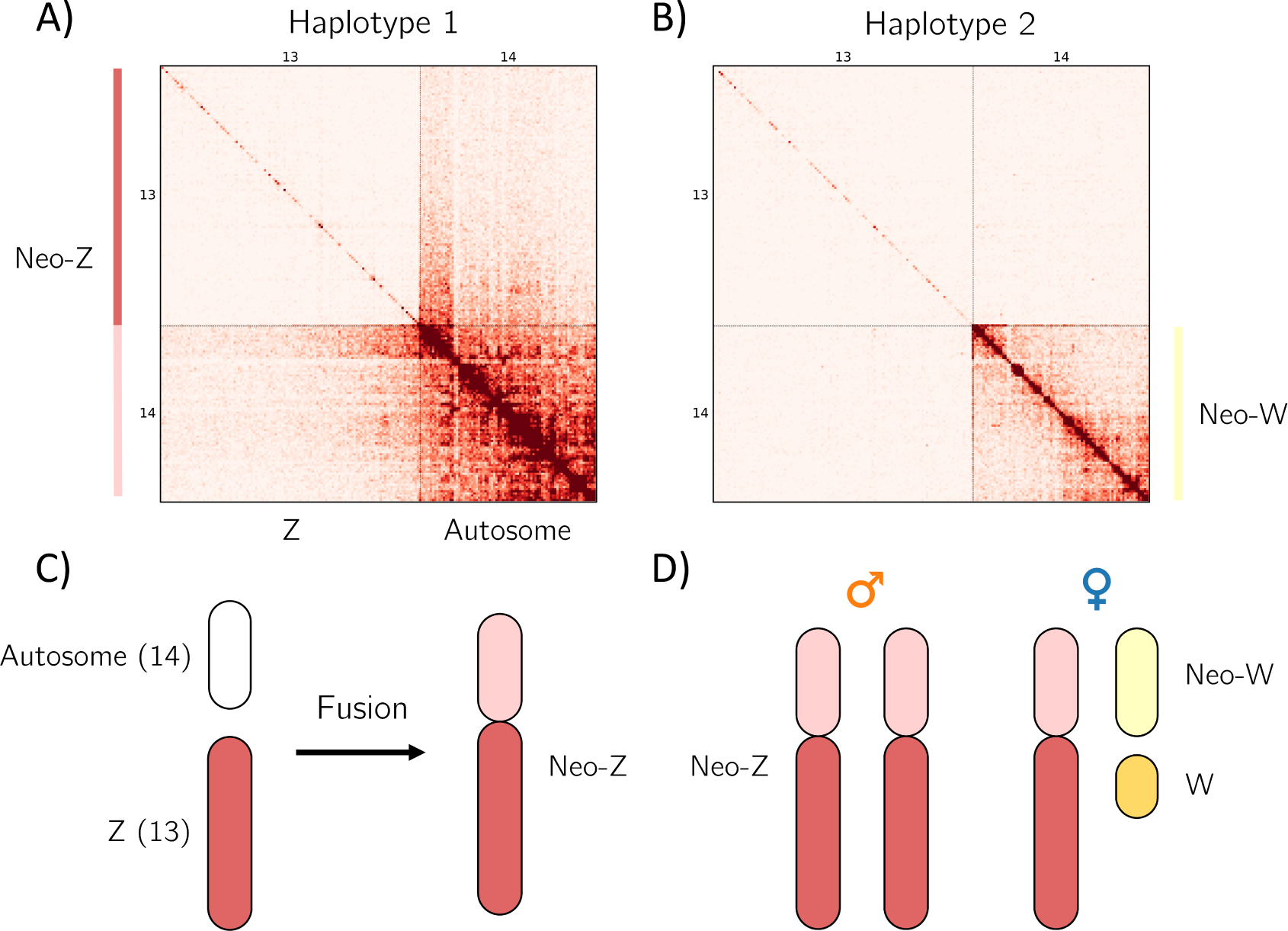
Z-autosome fusion in *M. ines*. The Hi-C map of chromosome 13 (Z) and chromosome 14 for the first haplotype (A) reveals inter-chromosomal contacts indicative of a Z-autosome fusion. These are absent in the second haplotype (B). Schematic of the fusion between chromosome 14 and the Z chromosome (C) and the neo-sex chromosome karyotype (D). When comparing the neo-Z and neo-W, we refer to the neo-Z as the segment homologous to the neo-W.

### Population structure

We generated WGS data for a total of 15 *M. ines* butterflies from Iberia and the Maghreb (Figure 2A, Table S1). A principal component analysis revealed three distinct population clusters: PC1 separated samples from the Iberian Peninsula and the Maghreb (51% of the variance explained), while PC2 separated Western and Eastern populations of the Maghreb (Figure 2B). Concordant with the PCA, substantial genetic structure was observed between North-African and European populations of *M. ines* (*F*_ST_ = 0.38, *d*_xy_ = 0.032), indicating a deep split between the continents. Western and Eastern populations of the Maghreb also showed evidence of population structure (*F*_ST_ = 0.20, *d*_xy_ = 0.025). Nucleotide diversity differed among the three populations, being highest in Eastern Maghreb (*π*_4D_ = 0.021), intermediate in Western Maghreb (*π*_4D_ = 0.013), and lowest in Iberia (*π*_4D_ = 0.008; Table 1).

**Figure 2.**
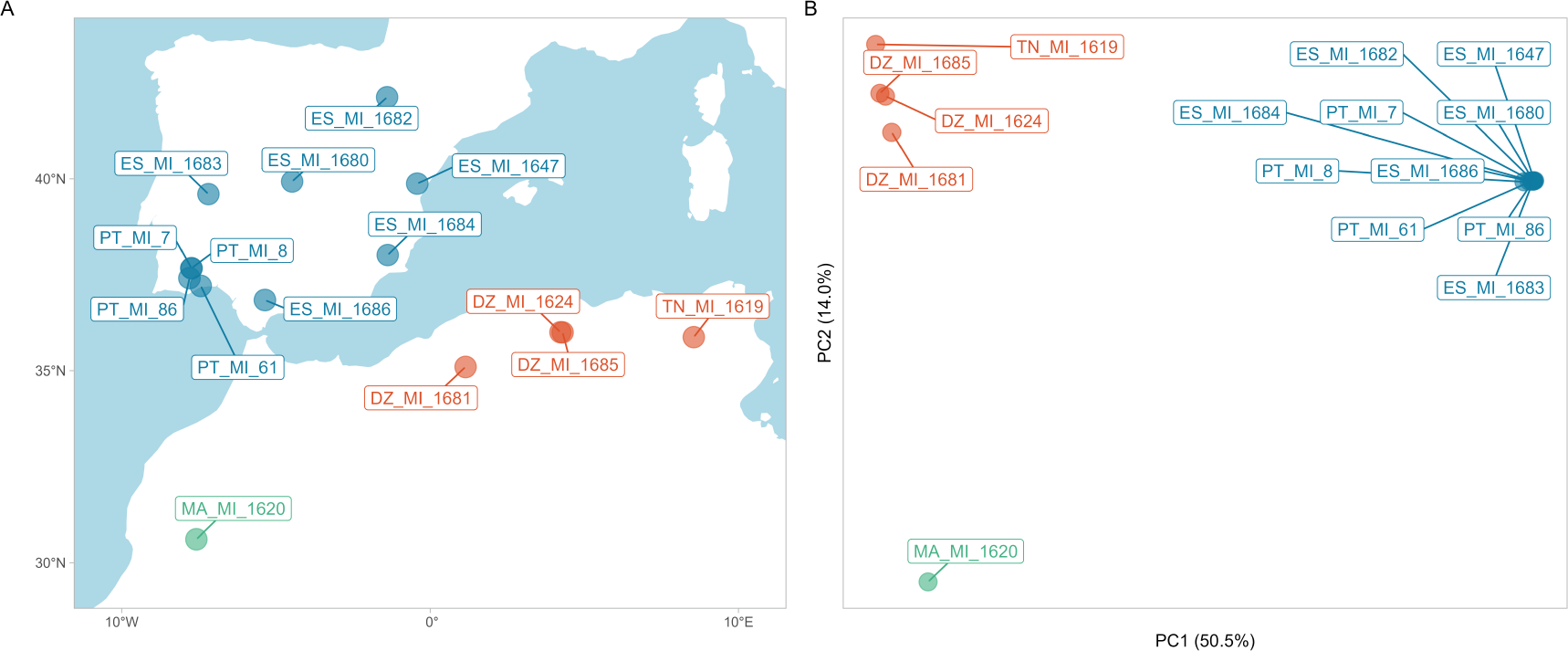
A) The *M. ines* WGS samples include ten individuals from Iberia, and five from the Maghreb. Samples are colored according to PCA clusters. B) Iberian, Western and Eastern Maghreb samples form three distinct clusters in a PCA: PC1 separates Iberia and North-Africa, PC2 separates Eastern and Western Maghreb samples.

**Table 1.** Nucleotide diversity at fourand zero-fold degenerate sites for autosomes and the neo-sex chromosome.

| Autosomes | $\pi_{4D}$ | $\pi_{0D}$ |
| --- | --- | --- |
| Iberia | 0.0075 | 0.0016 |
| Eastern Maghreb | 0.020 | 0.0032 |
| Western Maghreb | 0.013 | 0.0021 |
| Neo-sex chromosome |  |  |
| Neo-Z | 0.0072 | 0.0019 |
| Neo-W | 0.00077 | 0.00027 |
| Eastern Maghreb | 0.017 | 0.0033 |
| Western Maghreb | 0.012 | 0.0027 |

### Neo-sex chromosome distribution and frequency

The haplotype-specific HiC maps show the presence of a neo-Z chromosome in one female individual from Portugal. Large structural variants, such as chromosome fusions and inversions, cannot be directly detected with short-read sequencing data. However, the complete shut down of recombination between the neo-W and neo-Z will lead to predictable patterns in WGS data. First, the density of heterozygous sites on the neo-sex chromosome is expected to be significantly higher in heterogametic females (ZW) compared to homogametic males (ZZ). Second, with highly diverged, female-specific, neo-W chromosomes, the genetic structure for the neo-sex chromosome should reflect the sexual structure of the population. In other words, genetic variation is expected to cluster by sex for the neo-sex chromosome, with females being genetically closer to each other than to males, and *vice versa*. Neither pattern is expected for autosomes, which are recombining and not sex restricted.

For chromosome 14 Iberian females show a high density of heterozygous sites at fourfold degenerate (i.e. putatively neutral) positions of codons (*H*_4D_ = 0.027), while *H*_4D_ in males is significantly lower (0.006) and similar to autosomal *H*_4D_ (*∼* 0.006; Figure 3). This is consistent with Iberian females possessing diverged neo-Z and neo-W chromosomes (i.e. *H*_4D_ in females reflects divergence between the neo-Z and neo-W), whereas males possess two copies of the neo-Z chromosome.

**Figure 3.**
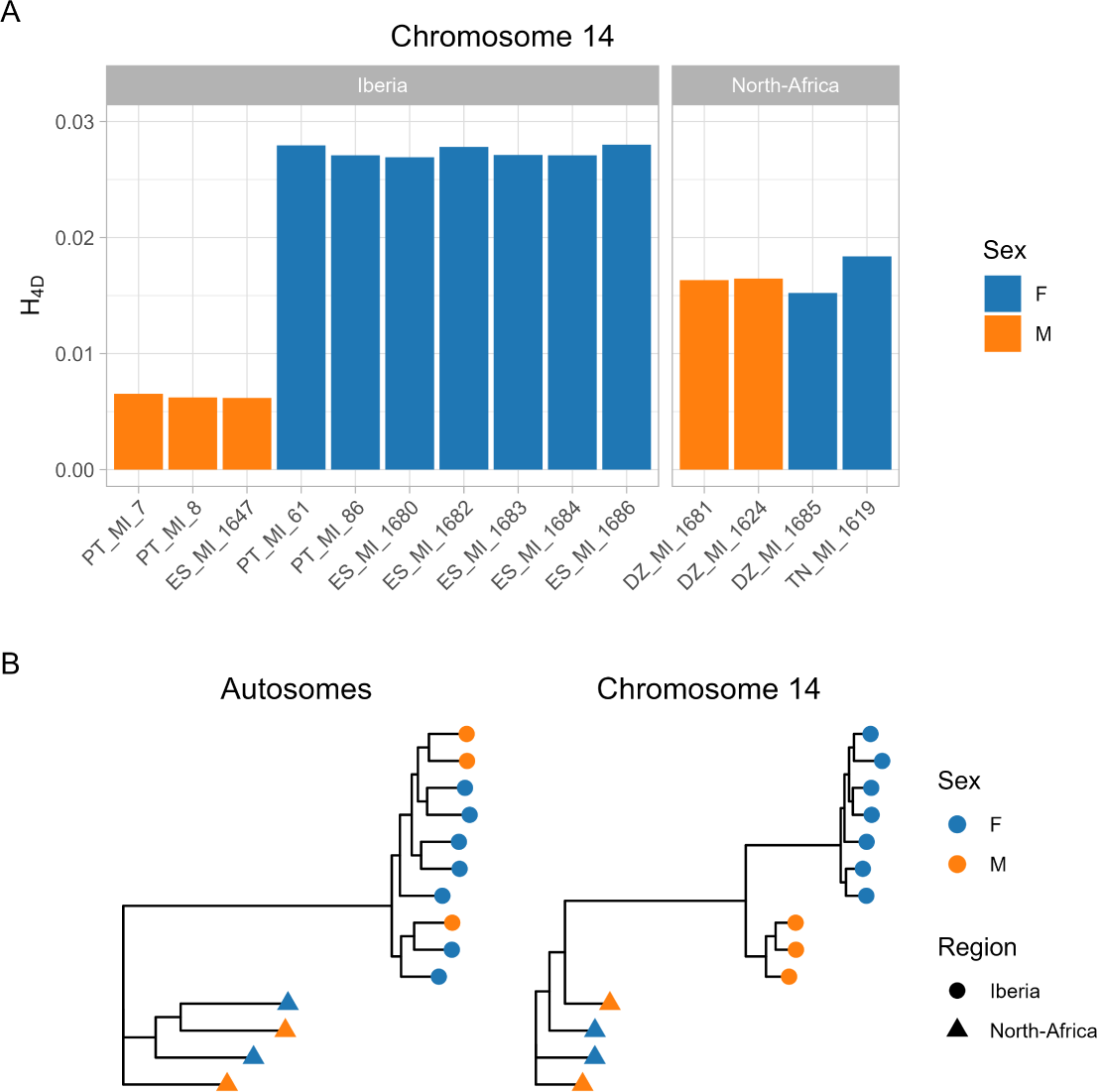
Detection of the neo-sex chromosome. A) Per-site heterozygosity (*H*_4D_) on chromosome 14 across Iberian and North-African samples is substantially larger in Iberian females than in Iberian males. By contrast, heterozygosity is similar across Eastern Maghreb samples, irrespective of sex. B) Maximum likelihood phylogenetic trees for the autosomes and chromosome 14. Clustering by sex is observed for chromosome 14, but only for Iberian samples.

A PCA of autosomal SNP data for Iberian samples shows that the genetic structure in Iberia reflects the underlying geographic structure of the sample (Figure S2). In contrast, we find strong clustering by sex on chromosome 14: PC1 separates samples by sex and explains 49.1% of the total variance (Figure S2). Similarly, a phylogenetic tree constructed from autosomal sequences reflects geography whereas the phylogeny for chromosome 14 is structured by sex (Figure 3). These patterns show that the neo-W chromosome is present in all Iberian females sampled. Assuming binomial sampling, the 95% confidence interval for the population frequency of the neo-W in Iberia is 0.59 *−* 1.

In contrast, in Eastern Maghreb samples, *H*_4D_ on chromosome 14 is similar for males and females. Additionally, genetic structure for chromosome 14 is similar to that of other autosomes and does not clusters by sex (Figure 3). These results lead us to conclude that the neo-sex chromosomes are absent from the Maghreb and therefore confined to Iberia.

### Age of the Z-autosome fusion

To determine whether the neo-sex chromosomes originated before or after the split between the Iberian and Maghreb populations, we modelled both the divergence history of *M. ines* populations and the neo-Z and the neo-W chromosomes using a blockwise site frequency spectrum approach implemented in gIMble (Laetsch et al. 2022). Given the genetic structure between Western and Eastern Maghreb samples, the single Western Maghreb individual was removed from the analysis. We fitted three different models to the autosomal data to infer the history of divergence between the Iberian and the Eastern Maghreb populations: i) Strict divergence between Iberia and Eastern Maghreb (*DIV*), i.e. no post-divergence gene flow, ii) an Isolation with migration with a constant rate of gene flow from Iberia to Eastern Maghreb (*IM*_→EM_) or in the opposite direction (*IM*_→IP_), iii) a Migration-only model, with migration in either direction (*MIG*_→EM_ or *MIG*_→IP_) but not involving a divergence of a common ancestral population into two.

The strict divergence model (*DIV*) best fit the data, i.e. *IM* models converge to the *DIV* model (*m_e_*= 0) and *MIG* models are less well supported (Table S2). Under the *DIV* model we infer a split between Eastern Maghreb and Iberia (Table 2) at time *T ≈* 1.5 *×* 10^6^ generations ago. Note that since *M. ines* is univoltine time estimates in generation and years are equivalent.

**Table 2.** Maximum composite likelihood estimate of the strict divergence model parameters for the three plateaus on the neo-sex chromosome, and the split between the Iberian Peninsula and Eastern Maghreb

| | $N_{eNeo-W}$ | $N_{eNeo-Z}$ | $N_{eAnc}$ | T | ln CL | $d_{xy}$ (intergenic) |
| --- | --- | --- | --- | --- | --- | --- |
| Plateau 1 | $2.1 \times 10^4$ | $2.7 \times 10^5$ | $1.3 \times 10^6$ | $1.4 \times 10^6$ | -958,297 | 0.022 |
| Plateau 2 | $2.1 \times 10^4$ | $2.6 \times 10^5$ | $9.8 \times 10^5$ | $8.6 \times 10^5$ | -130,194 | 0.015 |
| Plateau 3 | $2.1 \times 10^4$ | $2.4 \times 10^5$ | $5.7 \times 10^5$ | $5.0 \times 10^5$ | -31,886 | 0.008 |
| | $N_{eIberia}$ | $N_{eEastern\ Maghreb}$ | $N_{eAnc}$ | T | ln CL | $d_{xy}$ (intergenic) |
| Autosomes | $3.6 \times 10^5$ | $1.2 \times 10^6$ | $1.1 \times 10^6$ | $1.5 \times 10^6$ | -303,465,712 | 0.022 |

We also fitted a model of strict divergence between the neo-Z and the neo-W, excluding the last 2.5 Mb of the neo-sex chromosomes (see next section). The maximum composite likelihood population divergence time, which should reflect the onset of recombination arrest, was estimated at 1.4 *×* 10^6^ generations ago. This estimate for the origin of the neo-sex chromosomes is slightly more recent than the inferred split between Maghreb and Iberian populations. However, given the large confidence intervals of time estimates (in particular those for the neo-sex chromosomes divergence) we do not have power to determine whether the origin of the neo-sex chromosome pre- or postdated the split between Iberian and North-African populations (Figure S4)

### Plateaus of divergence suggest historical recombination events between the neo-Z and neo-W chromosomes

Since the neo-W is inherited as a single non-recombining haplotype, we would expect divergence between the neo-Z and the neo-W to be relatively uniform along the neo-sex chromosome (except for local variation in mutation rate and coalescence time). Surprisingly, a change point analysis of windowed divergence between neo-Z and neo-W sequences as measured by the density of heterozygous sites (*H*) in Iberian females revealed three distinct plateaus of divergence (Figure 4). The two plateaus exhibiting reduced divergence encompass the last 2.5 Mb of the neo-sex chromosome. These plateaus of neo-sex chromosome divergence were observed across all Iberian females, i.e. individual *H* landscapes all show the same three plateaus (Figure S3).

**Figure 4.**
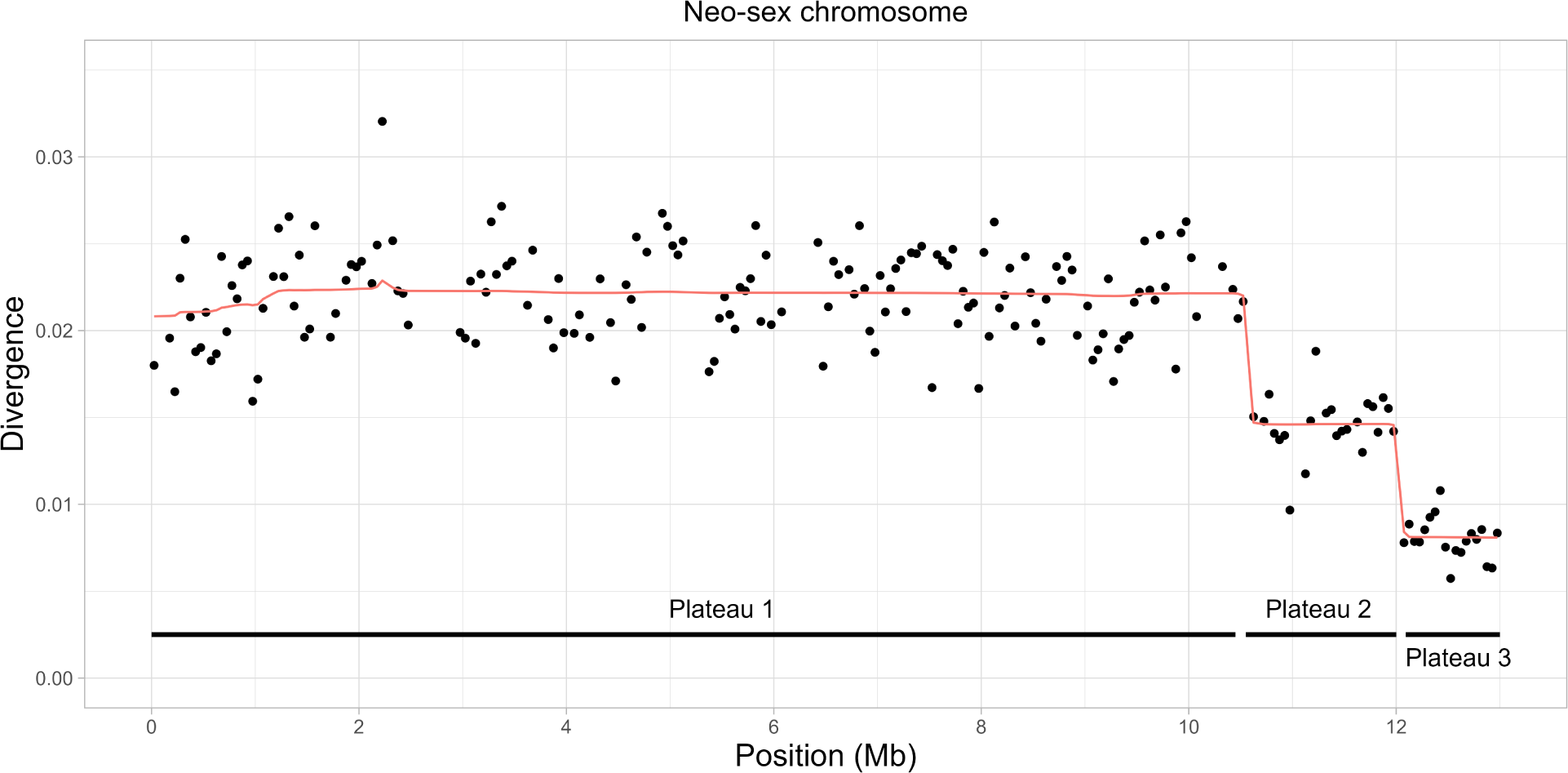
Divergence between the neo-sex chromosomes (as measured by *H*), showing three distinct plateaus. Reduced divergence between the neo-W and the neo-Z chromosome is observed for the last two plateaus (2.5 Mb at the end of the chromosome)

The largest plateau, which encompasses the first 10.5 Mb of the chromosome, shows the greatest divergence between neo-Z and neo-W sequences and likely reflects the timing of the Z-autosome fusion (see previous section). The last two plateaus show reduced divergence between the neosex chromosomes, suggesting a history of recombination between the neo-W and the neo-Z. Such a scenario is surprising given that meiosis in female Lepidoptera is achiasmatic. However, rare crossover events are the most plausible explanation given the scale of the plateaus (*>* 1 Mb), and the fact that they are separated by sharp boundaries.

Repeating the demographic analysis in the previous section, but focusing on plateaus 2 and 3, allows for an estimate of the timing of recombination events. The maximum composite likelihood estimate of *T* for plateaus 2 and 3 were 8.6 *×* 10^5^ and 5.0 *×* 10^5^ generations ago, respectively (Table 2).

### Patterns of sequence polymorphism on the neo-W

Given the evidence for recombination between the neo-Z and neo-W, we tested whether genetic diversity on the neo-W is consistent with a single tree, i.e. we asked whether there is any evidence for recent recombination events that are undetectable using windowed neo-W neo-Z divergence. Assuming that the neo-W chromosomes share a common ancestor more recently than the last recombination event between the neo-W and neo-Z, we expect neo-W variation to be consistent with a single genealogy. In line with this expectation, we find that 96% (3,240 out of 3,362) of the phased variants on the neo-W are compatible with a single tree, assuming an infinite sites mutation model and maximum parsimony. The 11 most common mutation types on the neo-W correspond to branches in the majority tree (Figure 5). Moreover, the small fraction of parsimony informative variants that are inconsistent with the majority topology do not share a particular topology and are not physically clustered (as would be expected for local genealogies generated by recombination events). This suggests that these are likely the result of back-mutations and genotyping errors, rather than recombination events.

**Figure 5.**
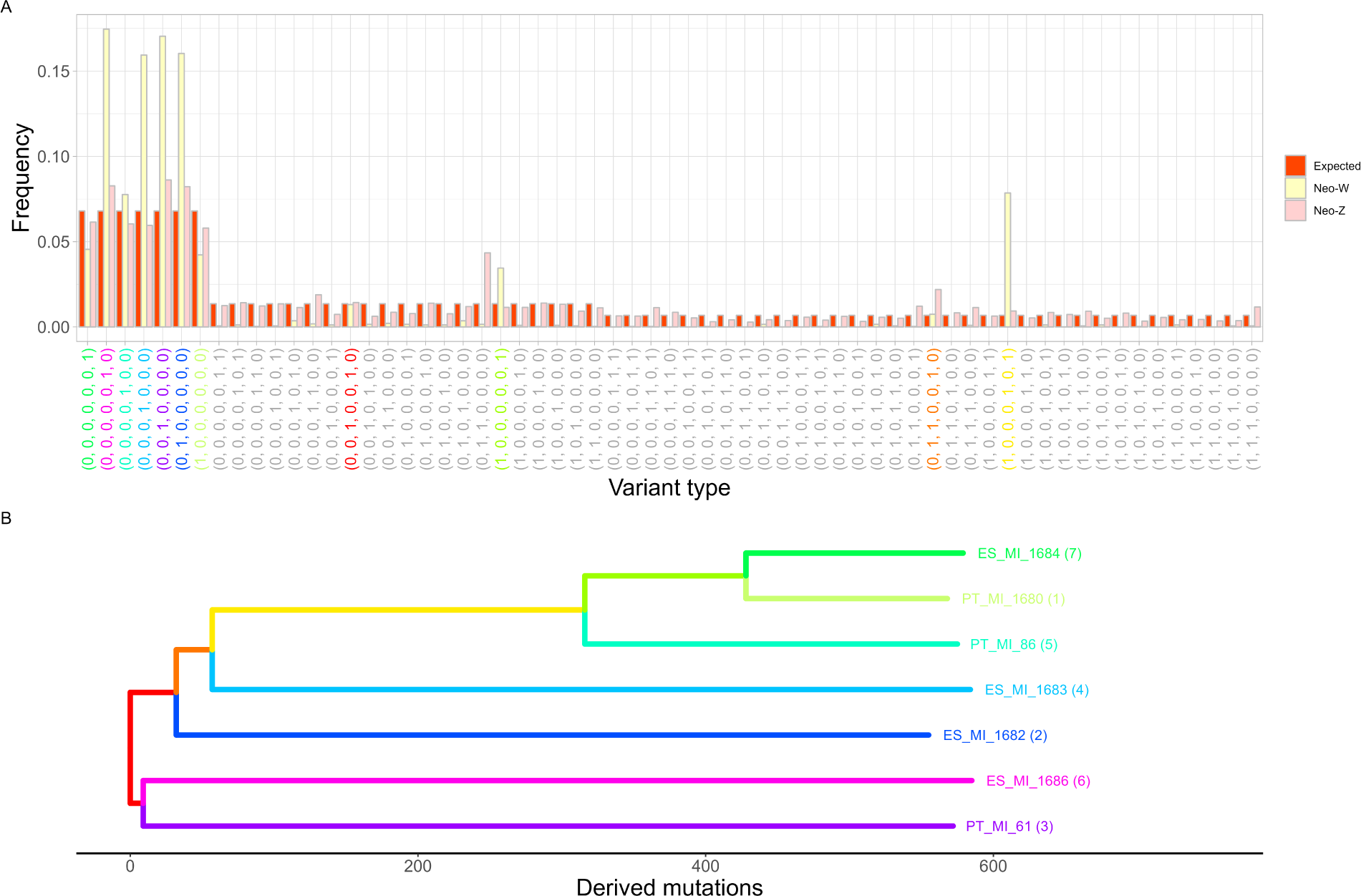
A) The folded variant type spectrum of the neo-Z and neo-W chromosomes and the corresponding expectation under the standard neutral coalescent. Variant types are coloured according to their corresponding branches in the neo-W tree shown in B).

The pattern of neo-W diversity which reflects a single tree is in stark contrast with the diversity among the (recombining) neo-Z haplotypes, which includes every possible type of variant. All models of population history that assume no population structure predict equal frequencies of all variants in a particular SFS class. We find that the mutation spectrum of the neo-Z haplotypes sampled from the seven Iberian females fits this simple expectation surprisingly well (Figure 5).

In the absence of natural selection and male biased mutation rate, and assuming an equal sex ratio, neo-W diversity is expected to be <u>^1^</u> of autosomal diversity. We find that neutral site diversity among *M. ines* neo-W chromosomes is extremely low (*π*_4D_ = 0.0008) and only *∼* <u>^1^</u> ^th^ of autosomal diversity (*π*_4D_ = 0.008). This result is unsurprising given that natural selection will remove diversity across the entire length of this non-recombining chromosome. Interestingly, the high proportion of singleton mutations among neo-W chromosomes (corresponding to external branches; Figure 5 B) suggests a past selective sweep. It is, however, difficult to glean the selective history of a non-recombining chromosome, given that there is only a single genealogy (see Discussion).

Given the reduced mutation load across plateau 2 and 3 (see next section and Figure 6) we hypothesised that recombinant neo-W haplotypes were favoured by natural selection. However, given the lack of recombination on the neo-W, the information about its selective history is limited and contained entirely within the neo-W genealogy (Figure 5). Thus, it would be inappropriate to employ genome-scan methods for inferring selective sweeps. We find that the *T*_MRCA_ of the neo-W tree (*∼* 5 *×* 10^4^) is an order of magnitude younger than *T*_Plateau 3_ (5.0 *×* 10^5^). Thus, any signal of selective sweeps occurring *>* 5.0 *×* 10^4^ generations ago has since been lost.

**Figure 6.**
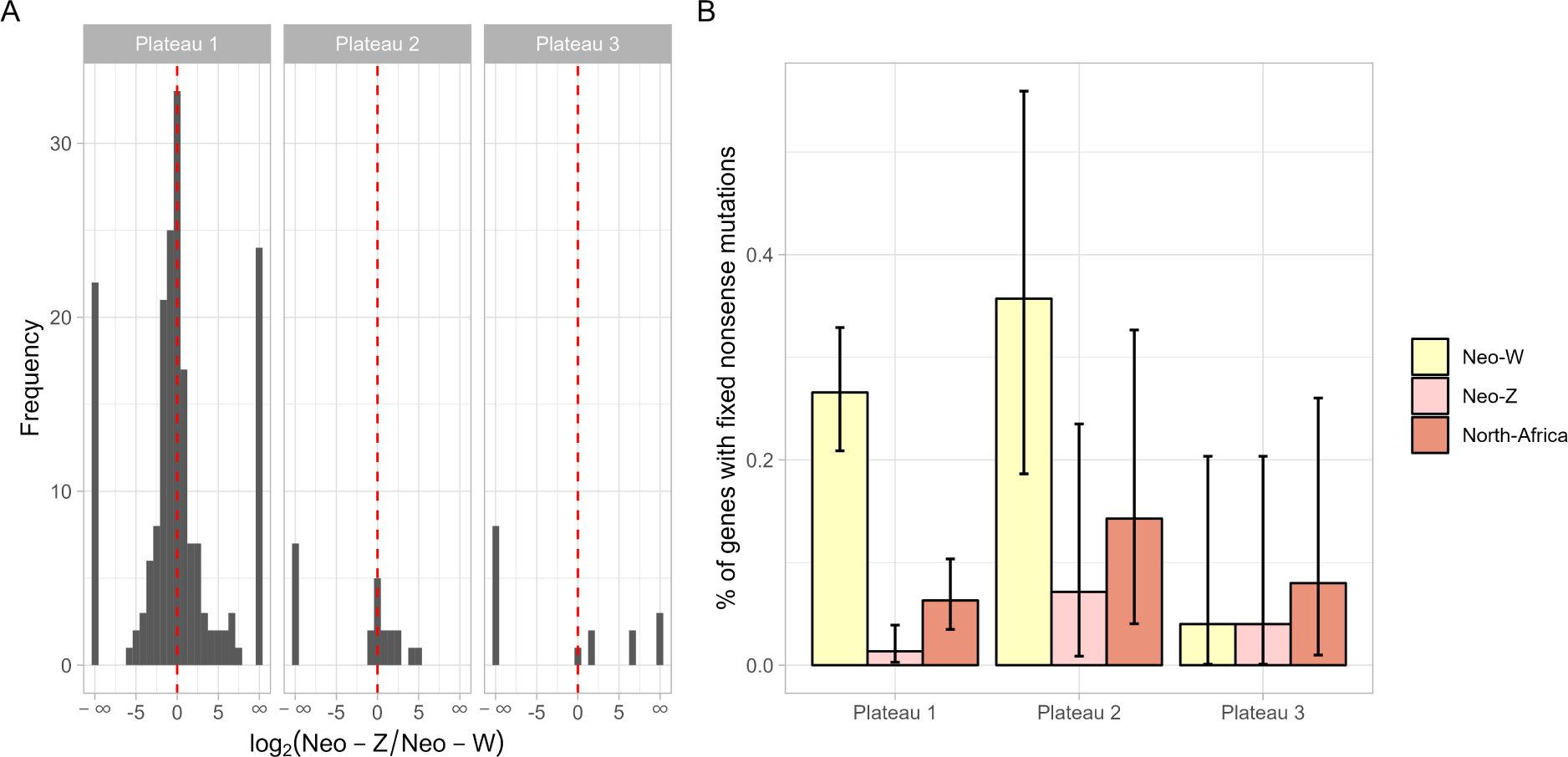
Neo-W chromosome degeneration. A) Gene expression of the neo-Z and neo-W chromosome, normalised by total expression (neo-Z + neo-W). B) The proportion of genes with fixed loss-of-function mutations on the neo-W is significantly greater for the oldest plateau (1) compared to both the neo-Z and the youngest plateau (3). Bars represent 95% binomial confidence intervals.

### Neo-W degeneration

The neo-W chromosome contains three plateaus that started degenerating at different time points, including one for which recombination arrest was recent (*∼* 0.5 Mya). This provides an opportunity to investigate the temporal dynamics of sequence degeneration due to recombination suppression. Here we quantify neo-W degeneration relative to homologous neo-Z sequence in terms of loss-of-function variants and gene expression.

We assessed sequence homology between the neo-Z and neo-W chromosome by comparing read depth in males and females. As expected, males were equally covered on the neo-sex chromosome and the autosomes (neo-sex:autosome depth ratio = 1). In contrast, the neo-sex chromosome in females showed reduced read depth (neo-sex:autosome depth ratio = 0.9) suggesting that only 0.8 neo-W reads map for every 1 neo-Z read. For plateau 1 neo-W chromosome reads had lower coverage than reads from the orthologous region of the Maghreb autosome (Figure S5, S6), despite having started diverging from the neo-Z at around the same time. This suggests that there has been sequence loss on the neo-W not experienced by the neo-Z or Maghreb autosome. By contrast, coverage did not significantly differ between the neo-Z and the neo-W on Plateau 3 (Wilcoxon rank-sum test, *p >* 0.4), consistent with limited divergence and sequence loss in this region.

We next considered the frequency of deleterious variants on the neo-W, neo-Z and the homologous Maghreb autosome. Consistent with substantial neo-W degeneration, 25% (95% binomial CI = 0.20 - 0.31) of genes contained loss-of-function mutations that were fixed on the neo-W, compared to 2% (0.008 - 0.05) on the neo-Z and 8% (0.04 - 0.11) on the Maghreb autosome. Across the neo-W, 27% (0.21 - 0.33) of genes within plateau 1 contained fixed loss-of-function mutations, compared to only 4% (0.001 - 0.20) of genes on plateau 3 (Figure 6). We also find that the neo-W has an elevated ratio of nucleotide diversity between 0D and 4D sites (*π*_0_*/π*_4_ = 0.35), compared to the neo-Z (0.26) and the Maghreb autosome (0.20). Elevated *π*_0_*/π*_4_ is consistent with a reduced efficacy of selection and an accumulation of weakly deleterious mutations at 0D sites.

We used previously published RNA-seq data from a single female (Mackintosh et al. 2019) to analyse haplotype-specific gene expression on the neo-sex chromosomes. Perhaps surprisingly, we do not find evidence of chromosome-wide gene silencing on the neo-W chromosome. Moreover, we find that the expression of neo-W genes is independent of degeneration. The expression level of genes with and without loss-of-function mutations were not statistically different (Wilcoxon rank-sum test, *p >* 0.1; Figure S7).

Overall, we find that the neo-W has lost DNA, and premature stop-codons and frame-shift mutations have become fixed. Yet, neo-W and neo-Z chromosomes have similar levels of expression, even at genes with loss-of-function mutations.

## Discussion

### A young neo-sex chromosome system

Neo-sex chromosomes provide a means of investigating the evolutionary consequences of recombination arrest. Here we have focused on a neo-sex chromosome system in Iberian *Melanargia ines* butterflies, where a recent Z-autosome fusion has resulted in the homologous autosome becoming a non-recombining neo-W chromosome. Our analysis of genome sequence data suggests that the neo-sex chromosomes originated *∼* 1.4 *×* 10^6^ generations ago. This neo-W is therefore older than the neo-Y of *D. albomicans* — the youngest neo-Y identified among *Drosophila* species (Zhou et al. 2012) — but younger than the neo-Y of *D. miranada* (Bachtrog & Charlesworth 2002). Similar to *Drosophila* neo-Y chromosomes, the *M. ines* neo-W shows evidence of degeneration. For example, 32% of genes contain loss-of-function mutations on plateau 1 of the neo-W. This level of degeneration is greater than that observed on the *D. albomicans* neo-Y chromosome (only 2% of genes contain loss-of-function mutations, Zhou et al. 2012) but similar to what is observed on the *D. miranda* neo-Y (Bachtrog 2005). We do not find evidence of gene silencing on the neo-W despite extensive gene decay. Our results are therefore not consistent with models of degeneration by regulatory evolution whereby silencing predates degeneration. Without a *de novo* assembled neo-W it is challenging to accurately quantify sequence degeneration, as many TEs insertions or other deleterious structural variants are undetectable with mapping-based analyses. Nonetheless, the majority of genes on the neo-W lack loss-of-function mutations consistent with a young neo-sex chromosome system at an intermediate level of sequence degeneration.

### The evolutionary history of the neo-sex chromosomes

Our analysis suggests that the Z-autosome fusion and associated neo-sex chromosomes are restricted to the Iberian population of *M. ines*. While the Iberian Peninsula and the Maghreb were connected by land (Corńee et al. 2016) prior to the Zanclean flood *≈* 5.3 Mya, our demographic analyses suggest that the split between Iberian and North-African *M. ines* populations is much younger than the Strait of Gibraltar (1.5 Mya). This estimate of population divergence is based on intergenic sites, which do not evolve under strict neutrality in *M. ines* (*π*_4_*_D_ > π_intergenic_*) and – as a result – is likely an underestimate. Moreover, we calibrated the estimate with the mutation rate of *H. melpomene* — whose most recent common ancestor with *M. ines* lived *∼* 60 *−* 80 Mya (Kumar et al. 2017). However, despite these caveats the most likely scenario is that Iberian populations of *M. ines* were established long after the Zanclean flood.

Our estimates for the origin of the neo-sex chromosomes show that population and neo-sex chromosome divergence began at a similar time. However, given the overlapping confidence intervals of divergence times we are unable to infer a specific order for the two events. This task is especially difficult given that coalescent simulations do not yet accommodate population-specific recombination rates. We instead assumed recombination on the neo-W, and, as a result, the estimated confidence interval for the neo-sex chromosome are likely too narrow. Additionally, the comparison of split times assumes that the autosomes and sex chromosomes have equal mutation rates. We do not know if it is the case in *M. ines*, or Lepidoptera in general. We note that male biased mutation rates have been observed in many organisms including *D. melanogaster* (Wang et al. 2023).

Although we have been unable to infer the full history of the neo-sex chromosomes, a plausible scenario is that the Iberian population was established from an ancestral North African population by migration, with the neo-sex chromosome arising and fixing shortly after in the isolated Iberian population. However, PSMC trajectories of *N_e_*change suggest no Iberian specific signal of an Out-of-Africa bottleneck (Fig S8).

### Historical recombination events

We find evidence for two historical recombination events between the neo-Z and the neo-W. One possibility is that recombination occurred in females. Rare spontaneous recombination despite achiasmy — both chiasma formation during meiosis and pre-meiotic ectopic recombination — has been observed in males of several Drosophila species: *D. melanogaster*, *D. simulans*, *D. virilis*, *inter alia* (Woodruff & Thompson 1977). Alternatively, rare migration into Iberia from populations with the ancestral (unfused) chromosome arrangement could have led to the neo-W migrating into males. The F1 offspring of a migrant and an Iberian individual will include individuals with a W chromosome, a neo-W, a Z and an unfused autosome. In these F1s, the neo-W and the W are not linked, allowing the neo-W to be transferred to males by independent assortment. Recombination between the neo-Z and the neo-W could therefore have happened in heterokaryotypic males. Although we do not find direct evidence for this scenario in *M. ines*, it is an expected outcome of rare migration events. A similar scenario has been reported in *D. americana*; individuals with a derived X-autosome fusion reproduce frequently with those carrying the ancestral arrangement, thereby allowing recombination between the neo-X and the neo-Y in heterokaryotypic males (Charlesworth et al. 1997, McAllister 2003). While our results are compatible with rare crossover events in female Lepidoptera (which has never been detected), we stress that the population genomic signal of rare female recombination is indistinguishable from recombination in (presumably also rare) descendants of migrant individuals, i.e. we cannot discern between these two scenarios. How plausible the male escape hypothesis is depends on the strength of postzygotic and premating barriers between African and Iberian *M. ines* populations. Differences in wing patterns and habitat between populations on either side of the Strait of Gibraltar have led some taxonomists to consider North African *M. ines* a distinct form (Wagner 1913).

Note that the rare recombination between sex chromosomes in achiasmatic taxa we have found here can be viewed as the inverse of strata formation via rare recombination suppressors in classical models of sex chromosome evolution (in taxa where both sexes recombine) (Charlesworth 1978). We have therefore chosen the term “plateau” to highlight the distinction from strata.

Our analysis of sequence degeneration suggests that these historical recombination events have locally reduced the mutation load on the neo-W by generating a haplotype where one portion (recently derived from the neo-Z chromosome) is free from the load associated with recombination arrest. We would expect such recombinant neo-W haplotypes to be under strong positive selection and to rapidly become fixed in the population. Importantly, there is a trade off between selection against neo-W load and potential neo-W linked female beneficial mutations. Thus, a recombinant neo-W haplotype may not always be beneficial, because female-specific alleles could be lost. However, the reduced mutation load across the youngest plateau 3 (Figure 6) gives reason to suspect that the recombinant neo-W haplotype swept through the population.

The neo-Z neo-W divergence time estimate for the 3rd plateau is an estimate of the time of the sweep that established the last recombinant neo-W/neo-W haplotype in the Iberian *M. ines* population. Conversely, the *T*_MRCA_ of the neo-W genealogy is a lower bound of the sweep time. However, since the neo W *T*_MRCA_ is an order of magnitude younger than the divergence of the 3rd plateau, neo-W diversity cannot contain any information about the sweep associated with the 3rd plateau. This is true even if one were to assume an extreme null model which assumes that the neo-W diversity evolves neutrally following the sweep (i.e. no background selection) and so recovers to 1/3 of that of the neo-Z. The strongly reduced genetic diversity on the neo-W (and our corresponding estimates of *N_W_*) suggest that – unsurprisingly – other strong selection must have been acting on the neo-W. Importantly, this will include selection on the mitochondrion, the ancestral W and any maternally inherited endosymbiont, given the maternal inheritance of the neo-W and its complete linkage to these other chromosomes. We find no evidence of any recent recombination affecting current neo-W diversity. While the fact that neo-W diversity can be placed on a single genealogy (Figure 3) simplifies analysis, it also means that there is limited power to make detailed inferences about the population processes that generated the neo-W genealogy.

Our finding of rare recombination between neo-sex chromosomes shows that sequence degeneration following recombination arrest can be a non-monotonic process, i.e. reversible, and mirrors previous results in certain *Drosophila* species (Charlesworth et al. 1997, McAllister 2003, Wei & Bachtrog 2019). As more neo-sex chromosomes are reported in Lepidoptera (Martin et al. 2020, Mackintosh et al. 2022, Rueda-M et al. 2023, Hööok et al. 2023), it will be interesting to see whether the neo-W degeneration and recombination observed in *M. ines* is representative of neo-sex chromosome evolution in this order.

## Materials and methods

### Sampling and sequencing

Fifteen butterflies were collected in Iberia and the Maghreb (Table S1). A high molecular weight (HMW) DNA extraction was performed for the genomic reference sample PT MI 8 using a salting out method (see Mackintosh et al. 2022 for details). For the other fourteen samples, DNA was extracted from thoracic tissue with a Qiagen DNeasy Blood & Tissue kit. DNA libraries were prepared with the Illumina Truseq Nano kit. The paired-end libraries were sequenced on an Illumina NovaSeq6000 machine. A Pacbio CLR library was prepared from the HMW DNA and sequenced on a Sequel I machine. Additionally, an Arima HiC reaction was performed with flash frozen thoracic tissue from sample PT MI 86. A TruSeq library was prepared from the crosslinked DNA and sequenced on an Illumina NovaSeq 6000.

### Genome assembly and haplotype-specific HiC maps

We generated a reference genome for *Melanargia ines* by assembling Pacbio continuous long reads with NextDenovo version 2.4.0 (Hu et al. 2023). The contig sequences were polished with Illumina short-reads from the same individual (PT MI 8) using Hapo-G version 1.1 (Aury & Istace 2021). We identified and removed haplotypic duplicates and contigs deriving from other organisms using purge dups version 1.2.5 (Guan et al. 2020) and blobtools version 1.1.1 (Laetsch & Blaxter 2017) respectively. We mapped HiC data (from PT MI 86) to the contigs with bwa-mem version 0.7.17 (Li & Durbin 2009) and then used YaHS version 1.1a.2 and juicebox version 1.11.08 to scaffold the assembly into chromosome-level sequences (Zhou et al. 2023, Robinson et al. 2018). We also generated haplotype-specific HiC maps following Mackintosh et al. (2022) to further investigate the karyotype of the female individual used for HiC sequencing (PT MI 86).

### Gene annotation

Repetitive elements in the genome assembly were masked with Red (Girgis 2015). Two previously published RNA-seq datasets (Mackintosh et al. 2019) were aligned to the assembly with HISAT2 2.1.0 (Kim et al. 2019). The repeat masked genome assembly and RNA-seq alignments were used to annotate genes with braker2.1.5 (Stanke et al. 2006, 2008, Li et al. 2009, Barnett et al. 2011, Lomsadze et al. 2014, Buchfink et al. 2015, Hoff et al. 2015, 2019).

### Variant calling and filtering

Whole genome sequencing (WGS) reads were trimmed with fastp 0.20.0 (Chen et al. 2018) and aligned to the reference genome using bwa-mem (bwa 0.7.17; Li & Durbin 2009). Duplicate reads were marked with Sambamba 0.6.6 (Tarasov et al. 2015). Freebayes v1.3.2-dirty (Richter et al. 2020) was used to call variants.

We used the gIMble prep module (Laetsch et al. 2022) to filter variants: variants were normalised and decomposed with bcftools 1.12 (Danecek et al. 2021) and vcfallelicprimitives (Garrison et al. 2022), respectively. Single nucleotide polymorphisms (SNPs) with support from a single strand, or from unbalanced reads (present solely on the right or the left side of the alternate allele) or within 2 bp of non-SNP variants were excluded. Only SNPs with a minimum genotype quality of 10 and read depth between 8 and 3 *×* mean genome depth were retained. Callable regions of the genomes — i.e. with read depth between 8 and 3 *×* mean genome coverage — were identified using mosdepth 0.3.2 (Pedersen & Quinlan 2018). Excluded SNPs were removed from callable regions. A VCF containing indels was produced by filtering variants as described above, with the exception that indels were retained.

### Estimation of diversity and divergence

Mean and windowed estimates of genetic diversity (*π* and *H*), divergence (*d*_xy_), and differentiation (*F*_ST_ and *d_a_*) were computed with custom-made python scripts (see Data accessibility). Coding sites — four-fold (4D) and zero-fold (0D) — were classified with codingSiteTypes.py available at: https://github.com/simonhmartin/genomics_general. The neo-W chromosome has regions that map poorly to the neo-Z reference assembly leading to biased estimates of divergence. When computing neo-W neo-Z divergence we therefore removed the 20% of windows with the lowest neo-W coverage, as the neo-W was *∼* 80% covered. Bayesian change point detection for windowed estimates of diversity and divergence was conducted with the bcp package (Erdman & Emerson 2008).

### Neo-sex chromosome detection

The genetic structure of the neo-sex chromosome and the autosomes were characterised with (a) principal component analyses, and (b) phylogenetic tree constructions. For PCAs, eigenvectors and eigenvalues of the genotype matrices were computed with Scikit-allel v1.3.5 (Miles et al. 2021). For phylogenetic trees, IUPAC consensus sequences were created for each individual using bcftools with the consensus command. Consensus sequences were aligned with MAFFT v7.520 (Katoh et al. 2002) and trimmed with trimAl v1.4 (Capella-Gutíerrez et al. 2009). Maximum likelihood phylogenetic trees were inferred from trimmed alignments with IQ-TREE 2 (Minh et al. 2020).

### Neo-sex chromosome phasing

Partial phasing of variants on the neo-sex chromosomes was achieved by identifying neo-W-specific alleles and extracting reads containing such alleles, as devised by Martin et al. (2020). We first identified neo-W specific alleles as alternative alleles that are present in a single copy in each Iberian female but absent in all Iberian males using a custom python script. Aligned WGS reads containing neo-W-specific alleles were isolated using an adapted version of the script filterSAMbyTargetBase.py available at: https://github.com/simonhmartin/genomics_general. Variants were called with freebayes v1.3.2 and filtered using gIMble prep with the aforementioned parameters (except with the minimum read depth relaxed to 3, given the haploid nature of neo-W chromosomes).

### Demographic history inference

Demographic inference was performed with gIMble (Laetsch et al. 2022) which fits a model of isolation with migration (*IM*) assuming an ancestral population that splits into two derived populations at time *T* followed by unidirectional gene flow from one population to the other at rate *m_e_*. The effective population size *N_e_* of the ancestral and derived populations are allowed to differ. Two nested models assuming either strict divergence (*DIV*, *m_e_* = 0) or long-term migration (*MIG*, *T → ∞*) can also be fitted with gIMble.

We fitted *DIV*, *IM* and *MIG* models to autosomal sequence of *M. ines* to infer the demographic history of the Iberian and Eastern Maghreb populations. To infer the history of the neo-sex chromosome, we created pseudo-diploid neo-W data (by combining phased neo-W haplotypes). We fitted a *DIV* model between the neo-W pseudo-diploids and Iberian males (i.e. neo-Z diploids) with gIMble. In both cases, the analysis was restricted to intergenic sites, with a block size of 64 bp. We assumed a mutation rate of 2.9 *×* 10^−9^ mutations per site per generation, the estimated rate for *Heliconius melpomene* (Keightley et al. 2015). We assumed a generation time of one year as *M. ines* is univoltine.

### Neo-sex chromosome coverage

Mean read depth per base (coverage) was evaluated with mosdepth 0.3.2 (Pedersen & Quinlan 2018). We used the difference in coverage between males (neo-Z + neo-Z) and females (neo-W + neo-Z) to estimate the coverage of neo-W chromosomes relative to neo-Z chromosomes. More specifically, we estimated neo-W coverage as cov_W_ = ((cov_ZW_ *−* cov_ZZ_*/*2) *×* 2 = ((cov_Female_ *−* cov_Male_*/*2) *×* 2.

### Gene expression analysis

We used previously published RNA-seq data from a female (PT MI 61) (Mackintosh et al. 2019) to analyse gene expression on the neo-sex chromosome. In order to obtain haplotype-specific gene expression profiles on the neo-sex chromosome of the female PT MI 61, we identified neo-W and neo-Z diagnostic alleles from the WGS dataset. To minimise reference bias, we also created a pseudo-reference assembly with neo-W alleles from PT MI 61 (as in Wei & Bachtrog 2019). We then mapped the trimmed RNA-seq reads to both the reference assembly and pseuo-reference with HISAT2 2.1.0 (Kim et al. 2019). Neo-Z specific RNA-seq reads were extracted from the alignments to the reference assembly and neo-W specific RNA-seq reads were extracted from alignments to the pseudo-reference, as described the WGS reads. Gene expression was subsequently quantified with HTseq-count 2.0.2 (Anders et al. 2015).

### Non-functional genes

Loss-of-function mutations, here defined as premature stop codons, frame-shift mutations, stoploss and start-loss variants, were detected with SnpEff 5.1d (Cingolani et al. 2012) from the VCF containing both SNP and indel calls. We estimated the proportion of genes with fixed derived loss-of-function mutations on the neo-W, the neo-Z and the African homolog. We only considered genes with non-zero expression in the Iberian female PT MI 61, i.e. which were expressed either by the neo-Z or the neo-W chromosome. This excluded spurious genes from the analysis, as well as genes which could not be phased.

### Tree test

Given that the neo-W is a non-recombining chromosome, genetic variation among these sequences should reflect a single genealogy. For a single neo-W tree with seven tips, we expect 11 folded variant types. By contrast, we expect recombining sequences, such as the neo-Z, to show evidence of all 63 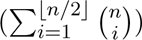 possible branches types (Felsenstein 1978). With this in mind, we summarised the neo-W and neo-Z haplotype alignments by extracting and counting SNP types and compared them to a neutral expectation.

We used the frequency of neo-W SNP types to construct an unrooted maximum parsimony tree. We estimated the root node age of the neo-W tree (time to the most common ancestor, *T*_MRCA_) as the average proportion of derived mutations (excluding fixed derived sites) per individual divided by the mutation rate. We polarised neo-W alleles with North-African alleles, using a custom python script. SNPs that were not consistent with the neo-W tree were discarded, whereas those which were consistent with the single tree but mispolarised [e.g. (0, 0, 0, 1) mispolarised as (1, 1, 1, 0)], were reassigned to the correct SNP type.

## Data accessibility

All the scripts and commands used for this project are available at: https://github.com/thomdec/ neosex_melanargia. The *Melanargia ines* genome assembly and all raw sequencing data are available at the European Nucleotide Archive under project accession PRJEB71083.

## Acknowledgements

This work was supported by an ERC starting grant (ModelGenomLand) to KL which supported TD. AM was supported by an E4 PhD studentship from the Natural Environment Research Council UK (NERC) (NE/S007407/1), KL was supported by a NERC research fellowship (NE/L011522/1). RV was supported by Grant PID2022-139689NB-I00 funded by MCIN/AEI/ 10.13039/501100011033 and by ERDF A way of making Europe, and by Departament de Recerca i Universitats de la Generalitat de Catalunya grant 2021 SGR 00420. Samples were obtained under permits DGPFEN/SEN/avp 19 12 and INAGA 500201/24/2014/1652. We thank Alex Hayward, Mohamed Ait Hammou, Sylvain Cuvelier, Leonardo Dapporto, Vlad Dincā, Juan Herńandez Roldán, Joan Carles Hinojosa, Rebbas Khellaf, Miguel Ĺopez Munguira, Michel Tarrier, Raluca Vodā for contributing samples, Simon Martin for helpful discussions and Deborah and Brian Charlesworth for insightful comments on an earlier draft and Edinburgh Genomics for generating the HiC and Pacbio data.

## References

Akagi, T., Varkonyi-Gasic, E., Shirasawa, K., Catanach, A., Henry, I. M., Mertten, D., Datson, P., Masuda, K., Fujita, N., Kuwada, E. et al. (2023), ‘Recurrent neo-sex chromosome evolution in kiwifruit’, Nature Plants 9(3), 393–402.

Anders, S., Pyl, P. T. & Huber, W. (2015), ‘HTSeq—a Python framework to work with highthroughput sequencing data’, Bioinformatics 31(2), 166–169.

Aury, J.-M. & Istace, B. (2021), ‘Hapo-G, haplotype-aware polishing of genome assemblies with accurate reads’, NAR Genomics and Bioinformatics 3(2). lqab034.

Bachtrog, D. (2005), ‘Sex chromosome evolution: molecular aspects of y-chromosome degeneration in drosophila’, Genome research 15(10), 1393–1401.

Bachtrog, D. (2008), ‘The temporal dynamics of processes underlying y chromosome degeneration’, Genetics 179(3), 1513–1525.

Bachtrog, D. (2013), ‘Y-chromosome evolution: Emerging insights into processes of Y-chromosome degeneration’, Nature Reviews Genetics 14(2), 113–124.

Bachtrog, D. & Charlesworth, B. (2002), ‘Reduced adaptation of a non-recombining neo-y chromosome’, Nature 416(6878), 323–326.

Barnett, D. W., Garrison, E. K., Quinlan, A. R., Strömberg, M. P. & Marth, G. T. (2011), ‘BamTools: a C++ API and toolkit for analyzing and managing BAM files’, Bioinformatics 27(12), 1691–1692.

Baumdicker, F., Bisschop, G., Goldstein, D., Gower, G., Ragsdale, A. P., Tsambos, G., Zhu, S., Eldon, B., Ellerman, E. C., Galloway, J. G., Gladstein, A. L., Gorjanc, G., Guo, B., Jeffery, B., Kretzschumar, W. W., Lohse, K., Matschiner, M., Nelson, D., Pope, N. S., Quinto-Cortés, C. D., Rodrigues, M. F., Saunack, K., Sellinger, T., Thornton, K., van Kemenade, H., Wohns, A. W., Wong, Y., Gravel, S., Kern, A. D., Koskela, J., Ralph, P. L. & Kelleher, J. (2021), ‘Efficient ancestry and mutation simulation with msprime 1.0’, Genetics 220(3), iyab229.

Buchfink, B., Xie, C. & Huson, D. H. (2015), ‘Fast and sensitive protein alignment using diamond’, Nature Methods 12(1), 59–60.

Capella-Gutiérrez, S., Silla-Martínez, J. M. & Gabaldón, T. (2009), ‘trimAl: A tool for automated alignment trimming in large-scale phylogenetic analyses’, Bioinformatics 25(15), 1972–1973.

Charlesworth, B. (1978), ‘Model for evolution of y chromosomes and dosage compensation’, PNAS 77(11), 5618–5622.

Charlesworth, B. & Charlesworth, D. (2000), ‘The degeneration of y chromosomes’, Philosophical Transactions of the Royal Society of London. Series B: Biological Sciences 355(1403), 1563–1572.

Charlesworth, B., Charlesworth, D., Hnilicka, J., Yu, A. & Guttman, D. S. (1997), ‘Lack of Degeneration of Loci on the Neo-Y Chromosome of Drosophila americana americana’, Genetics 145(4), 989–1002.

Chen, S., Zhou, Y., Chen, Y. & Gu, J. (2018), ‘Fastp: An ultra-fast all-in-one FASTQ preprocessor’, Bioinformatics 34(17), i884–i890.

Choo, K. A. (1998), ‘Why is the centromere so cold?’, Genome research 8(2), 81–82.

Cingolani, P., Platts, A., Wang, L. L., Coon, M., Nguyen, T., Wang, L., Land, S. J., Lu, X. & Ruden, D. M. (2012), ‘A program for annotating and predicting the effects of single nucleotide polymorphisms, SnpEff’, Fly 6(2), 80–92.

Cornée, J.-J., Muñch, P., Achalhi, M., Merzeraud, G., Azdimousa, A., Quillévéré, F., Melintedobrinescu, M., Chaix, C., Moussa, A. B., Lofi, J., Séranne, M. & Moissette, P. (2016), ‘The messinian erosional surface and early pliocene reflooding in the alboran sea: New insights from the boudinar basin, morocco’, Sedimentary Geology 333, 115–129.

Danecek, P., Bonfield, J. K., Liddle, J., Marshall, J., Ohan, V., Pollard, M. O., Whitwham, A., Keane, T., McCarthy, S. A., Davies, R. M. & Li, H. (2021), ‘Twelve years of SAMtools and BCFtools’, GigaScience 10(2), giab008.

Dapporto, L., Menchetti, M., Vodă, R., Corbella, C., Cuvelier, S., Djemadi, I., Gascoigne-Pees, M., Hinojosa, J. C., Lam, N. T., Serracanta, M., Talavera, G., Dincă, V. & Vila, R. (2022), ‘The atlas of mitochondrial genetic diversity for western palaearctic butterflies’, Global Ecology and Biogeography 31(11), 2184–2190.

de Lesse, H. (1970), ‘Formules chromosomiques de quelques rhopalocères paĺearctiques [lep]’, Bulletin de la Socíté entomologique de France 75(7), 214–216.

Erdman, C. & Emerson, J. W. (2008), ‘Bcp: An R Package for Performing a Bayesian Analysis of Change Point Problems’, Journal of Statistical Software 23, 1–13.

Felsenstein, J. (1978), ‘The Number of Evolutionary Trees’, Systematic Biology 27(1), 27>–33.

Fisher, R. A. (1930), The genetical theory of natural selection, Oxford University Press.

Garrison, E., Kronenberg, Z. N., Dawson, E. T., Pedersen, B. S. & Prins, P. (2022), ‘A spectrum of free software tools for processing the VCF variant call format: Vcflib, bio-vcf, cyvcf2, hts-nim and slivar’, PLOS Computational Biology 18(5), e1009123.

Gil-Fernández, A., Saunders, P. A., Martín-Ruiz, M., Ribagorda, M., López-Jiménez, P., Jeffries, D. L., Parra, M. T., Viera, A., Rufas, J. S., Perrin, N., et al. (2020), ‘Meiosis reveals the early steps in the evolution of a neo-xy sex chromosome pair in the african pygmy mouse mus minutoides’, PLoS Genetics 16(11), e1008959.

Girgis, H. Z. (2015), ‘Red: an intelligent, rapid, accurate tool for detecting repeats de-novo on the genomic scale’, BMC bioinformatics 16(1), 1–19.

Guan, D., McCarthy, S. A., Wood, J., Howe, K., Wang, Y. & Durbin, R. (2020), ‘Identifying and removing haplotypic duplication in primary genome assemblies’, Bioinformatics 36(9), 2896– 2898.

Hill, W. & Robertson, A. (1968), ‘Linkage disequilibrium in finite populations’, Theoretical and applied genetics 38, 226–231.

Hoff, K., Lange, S., Lomsadze, A., Borodovsky, M. & Stanke, M. (2015), ‘BRAKER1: Unsupervised RNA-Seq-Based Genome Annotation with GeneMark-ET and AUGUSTUS’, Bioinformatics 32(5), 767–769.

Hoff, K., Lomsadze, A., Borodovsky, M. & Stanke, M. (2019), Gene Prediction: Methods and Protocols, Springer New York, chapter Whole-Genome Annotation with BRAKER, pp. 65–95.

Höök, L., Vila, R., Wiklund, C. & Backström, N. (2023), ‘Temporal dynamics of faster neo-Z evolution in butterflies’, bioRxiv pp. 2023–12.

Howell, E. C., Armstrong, S. J. & Filatov, D. A. (2009), ‘Evolution of neo-sex chromosomes in silene diclinis’, Genetics 182(4), 1109–1115.

Hu, J., Wang, Z., Sun, Z., Hu, B., Ayoola, A. O., Liang, F., Li, J., Sandoval, J. R., Cooper, D. N., Ye, K. et al. (2023), ‘An efficient error correction and accurate assembly tool for noisy long reads’, bioRxiv pp. 2023–03. Unpublished. doi: 10.1101/2023.03.09.531669.

Huang, Z., De O. Furo, I., Liu, J., Peona, V., Gomes, A. J., Cen, W., Huang, H., Zhang, Y., Chen, D., Xue, T., et al. (2022), ‘Recurrent chromosome reshuffling and the evolution of neo-sex chromosomes in parrots’, Nature communications 13(1), 944.

Katoh, K., Misawa, K., Kuma, K.-i. & Miyata, T. (2002), ‘MAFFT: A novel method for rapid multiple sequence alignment based on fast Fourier transform’, Nucleic Acids Research 30(14), 3059– 3066.

Keightley, P. D., Pinharanda, A., Ness, R. W., Simpson, F., Dasmahapatra, K. K., Mallet, J., Davey, J. W. & Jiggins, C. D. (2015), ‘Estimation of the Spontaneous Mutation Rate in Heliconius melpomene’, Molecular Biology and Evolution 32(1), 239–243.

Kim, D., Paggi, J. M., Park, C., Bennett, C. & Salzberg, S. L. (2019), ‘Graph-based genome alignment and genotyping with HISAT2 and HISAT-genotype’, Nature Biotechnology 37(8), 907– 915.

Kumar, S., Stecher, G., Suleski, M. & Hedges, S. B. (2017), ‘TimeTree: A Resource for Timelines, Timetrees, and Divergence Times’, Molecular Biology and Evolution 34(7), 1812–1819.

Laetsch, D. & Blaxter, M. (2017), ‘Blobtools: Interrogation of genome assemblies’, F1000Research 6(1287).

Laetsch, D. R., Bisschop, G., Martin, S. H., Aeschbacher, S., Setter, D. & Lohse, K. (2022), ‘Demographically explicit scans for barriers to gene flow using gIMble’.

Lenormand, T., Fyon, F., Sun, E. & Roze, D. (2020), ‘Sex chromosome degeneration by regulatory evolution’, Current Biology 30(15), 3001–3006.e5.

Li, H. & Durbin, R. (2009), ‘Fast and accurate short read alignment with Burrows–Wheeler transform’, Bioinformatics 25(14), 1754–1760.

Li, H., Handsaker, B., Wysoker, A., Fennell, T., Ruan, J., Homer, N., Marth, G., Abecasis, G., Durbin, R. & Subgroup, . G. P. D. P. (2009), ‘The Sequence Alignment/Map format and SAMtools’, Bioinformatics 25(16), 2078–2079.

Lomsadze, A., Burns, P. D. & Borodovsky, M. (2014), ‘Integration of mapped RNA-Seq reads into automatic training of eukaryotic gene finding algorithm’, Nucleic Acids Research 42(15), e119– e119.

Mackintosh, A., Laetsch, D. R., Baril, T., Foster, R. G., Dincă, V., Vila, R., Hayward, A. & Lohse, K. (2022), ‘The genome sequence of the lesser marbled fritillary, brenthis ino, and evidence for a segregating neo-z chromosome’, G3 12(6), jkac069.

Mackintosh, A., Laetsch, D. R., Hayward, A., Charlesworth, B., Waterfall, M., Vila, R. & Lohse, K. (2019), ‘The determinants of genetic diversity in butterflies’, Nature Communications 10(1), 3466.

Martin, S. H., Singh, K. S., Gordon, I. J., Omufwoko, K. S., Collins, S., Warren, I. A., Munby, H., Brattström, O., Traut, W., Martins, D. J., Smith, D. A. S., Jiggins, C. D., Bass, C. & ffrench-Constant, R. H. (2020), ‘Whole-chromosome hitchhiking driven by a male-killing endosymbiont’, PLOS Biology 18(2), e3000610.

McAllister, B. F. (2003), ‘Sequence Differentiation Associated With an Inversion on the Neo-X Chromosome of Drosophila americana’, Genetics 165(3), 1317–1328.

Miles, A., io bot, p., R M., Ralph, P., Harding, N., Pisupati, R., Rae, S. & Millar, T. (2021), ‘Cggh/scikit-allel: V1.3.3’, Zenodo.

Minh, B. Q., Schmidt, H. A., Chernomor, O., Schrempf, D., Woodhams, M. D., von Haeseler, A. & Lanfear, R. (2020), ‘IQ-TREE 2: New Models and Efficient Methods for Phylogenetic Inference in the Genomic Era’, Molecular Biology and Evolution 37(5), 1530–1534.

Mongue, A. J., Nguyen, P., Voleníková, A. & Walters, J. R. (2017), ‘Neo-sex Chromosomes in the Monarch Butterfly, Danaus plexippus’, G3 Genes—Genomes—Genetics 7(10), 3281–3294.

Muller, H. J. (1964), ‘The relation of recombination to mutational advance’, Mutation Research/-Fundamental and Molecular Mechanisms of Mutagenesis 1(1), 2–9.

Pedersen, B. S. & Quinlan, A. R. (2018), ‘Mosdepth: Quick coverage calculation for genomes and exomes’, Bioinformatics 34(5), 867–868.

Richter, F., Morton, S. U., Qi, H., Kitaygorodsky, A., Wang, J., Homsy, J., DePalma, S., Patel, N., Gelb, B. D., Seidman, J. G., Seidman, C. E. & Shen, Y. (2020), ‘Whole Genome De Novo Variant Identification with FreeBayes and Neural Network Approaches’.

Robinson, J. T., Turner, D., Durand, N. C., Thorvaldsdóttir, H., Mesirov, J. P. & Aiden, E. L. (2018), ‘Juicebox.js provides a cloud-based visualization system for Hi-C data’, Cell Systems 6(2), 256–258.e1.

Rueda-M, N., Jiggins, C. D., Pardo-Diaz, C., Montejo-Kovacevich, G., McMillan, W. O., McCarthy, S., Ready, J., Kozak, K. M., Arias, C. F., Durbin, R., Meier, J. I. & Salazar, C. (2023), ‘Three sequential sex chromosome – autosome fusions in heliconius butterflies’, bioRxiv. **URL:** https://www.biorxiv.org/content/early/2023/03/24/2023.03.06.531374

Sacchi, B., Humphries, Z., Kružlicová, J., Bodláková, M., Pyne, C., Choudhury, B., Gong, Y., Băcovský, V., Hobza, R., Barrett, S. C. & Wright, S. I. (2023), ‘Phased assembly of neo-sex chromosomes reveals extensive y degeneration and rapid genome evolution in rumex hastatulus’, bioRxiv. **URL:** https://www.biorxiv.org/content/early/2023/10/06/2023.09.26.559509

Smith, D. A., Gordon, I. J., Traut, W., Herren, J., Collins, S., Martins, D. J., Saitoti, K., Ireri, P. & Ffrench-Constant, R. (2016), ‘A neo-W chromosome in a tropical butterfly links colour pattern, male-killing, and speciation’, Proceedings of the Royal Society B: Biological Sciences 283(1835), 20160821.

Stanke, M., Diekhans, M., Baertsch, R. & Haussler, D. (2008), ‘Using native and syntenically mapped cDNA alignments to improve de novo gene finding’, Bioinformatics 24(5), 637–644.

Stanke, M., Schöffmann, O., Morgenstern, B. & Waack, S. (2006), ‘Gene prediction in eukaryotes with a generalized hidden markov model that uses hints from external sources’, BMC Bioinformatics 7(1), 62.

Tarasov, A., Vilella, A. J., Cuppen, E., Nijman, I. J. & Prins, P. (2015), ‘Sambamba: Fast processing of NGS alignment formats’, Bioinformatics 31(12), 2032–2034.

Wagner (1913), ‘Eine neue lokalform von *Melanargia ines* hoffm.’, Int. Ent. Zs 7(17), 111–112.

Wang, Y., McNeil, P., Abdulazeez, R., Pascual, M., Johnston, S. E., Keightley, P. D. & J., O. D. (2023), ‘Variation in mutation, recombination, and transposition rates in Drosophila melanogaster and Drosophila simulans’, Genome Research 33(10), 587–598.

Wei, K. H. & Bachtrog, D. (2019), ‘Ancestral male recombination in drosophila albomicans produced geographically restricted neo-y chromosome haplotypes varying in age and onset of decay’, PLoS Genetics 15(11), e1008502.

Woodruff, R. C. & Thompson, J. N. (1977), ‘An analysis of spontaneous recombination in drosophila melanogaster males’, Heredity 38(3), 291–307.

Wright, C. J., Stevens, L., Mackintosh, A., Lawniczak, M. & Blaxter, M. (2023), ‘Chromosome evolution in lepidoptera’, bioRxiv pp. 2023–05. Unpublished. doi: 10.1101/2023.05.12.540473.

Zhou, C., McCarthy, S. A. & Durbin, R. (2023), ‘Yahs: yet another hi-c scaffolding tool’, Bioinformatics 39(1), btac808.

Zhou, Q., Zhu, H.-m., Huang, Q.-f., Zhao, L., Zhang, G.-j., Roy, S. W., Vicoso, B., Xuan, Z.-l., Ruan, J., Zhang, Y., Zhao, R.-p., Ye, C., Zhang, X.-q., Wang, J., Wang, W. & Bachtrog, D. (2012), ‘Deciphering neo-sex and B chromosome evolution by the draft genome of Drosophila albomicans’, BMC Genomics 13(1), 109.

