## Supplementary material for "Rewinding the ratchet: rare recombination locally rescues neo-W degeneration and generates plateaus of sex-chromosome divergence"

### Supplementary text

#### Phasing

Partial phasing of the neo-sex chromosome was achieved by identifying and extracting neoZ- and neo-W-specific reads. Neo-W diagnostic alleles were defined as alternative alleles present in single copy in all seven females, while absent in males, at sites where all females are heterozygous. On the neo-sex chromosome, 69% of sites heterozygous across all females had neo-W specific alleles. One might expect alleles that are present in a single copy in all seven females, and absent in all three males, purely by chance. However, this distribution of genotypes is extremely rare on other autosomes: only 0.4% of sites heterozygous across all females on the autosomes. This negligible error rate suggest that robustness of our simple phasing method. On average, 36% of female reads mapping to the reference neo-W were classified as neo-W reads. Since female had an average read depth of 90%, the neo-W was expected to account for 40% of the reads on the neo-sex chromosome, assuming that the neo-Z was fully covered. We thus estimate that $0.36/0.40 = 90\%$ of the neo-W chromosome was successfully phased.

##### Parametric bootstrap

We estimated confidence intervals around the inferred split time $T$ by parametric bootstrap. We generated bootstrap replicates of the best-fitting model with the gIMble wrapper for msprime (Baumdicker et al. 2021), and computed the maximum composite likelihoods (MCLEs) parameters of the replicates. For the origin of the neo-sex chromosome, we simulated 100 bootstrap replicates under the best-fitting model and fitted the replicates to a $DIV$ model. We assumed a recombination rate of 1.9 cM/Mb, i.e. one crossover per tetrad per male meiosis for the neo-sex chromosome, assuming 50% female in the population ($\frac{100\ \text{cM}\times 1/4}{13\ \text{Mb}} = 1.9$ cM/Mb). Idem for the autosomal demographic history of *M. ines*, except with a recombination rate of 0.7 cM/Mb — the weighted mean of autosome lengths (34 Mb) is greater than the size of the neo-sex chromosome (13 Mb), explaining the lower recombination rate for autosomal regions.

### References

Baumdicker, F., Bisschop, G., Goldstein, D., Gower, G., Ragsdale, A. P., Tsambos, G., Zhu, S., Eldon, B., Ellerman, E. C., Galloway, J. G., Gladstein, A. L., Gorjanc, G., Guo, B., Jeffery, B., Kretzschumar, W. W., Lohse, K., Matschiner, M., Nelson, D., Pope, N. S., Quinto-Cortés, C. D.,

Rodrigues, M. F., Saunack, K., Sellinger, T., Thornton, K., van Kemenade, H., Wohns, A. W., Wong, Y., Gravel, S., Kern, A. D., Koskela, J., Ralph, P. L. & Kelleher, J. (2021), 'Efficient ancestry and mutation simulation with msprime 1.0', *Genetics* **220**(3), iyab229.

### Supplementary figures and tables

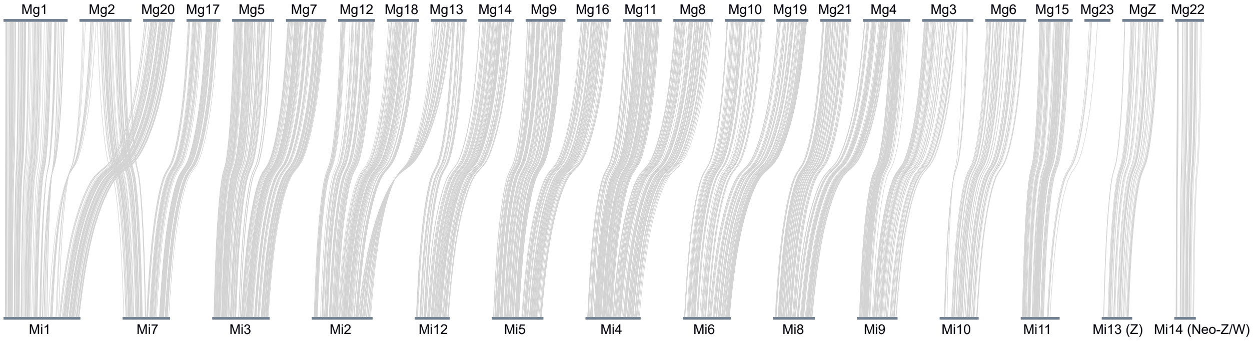


**Figure S1** – Minimap alignment between *M. ines* and *M. galathea* assemblies

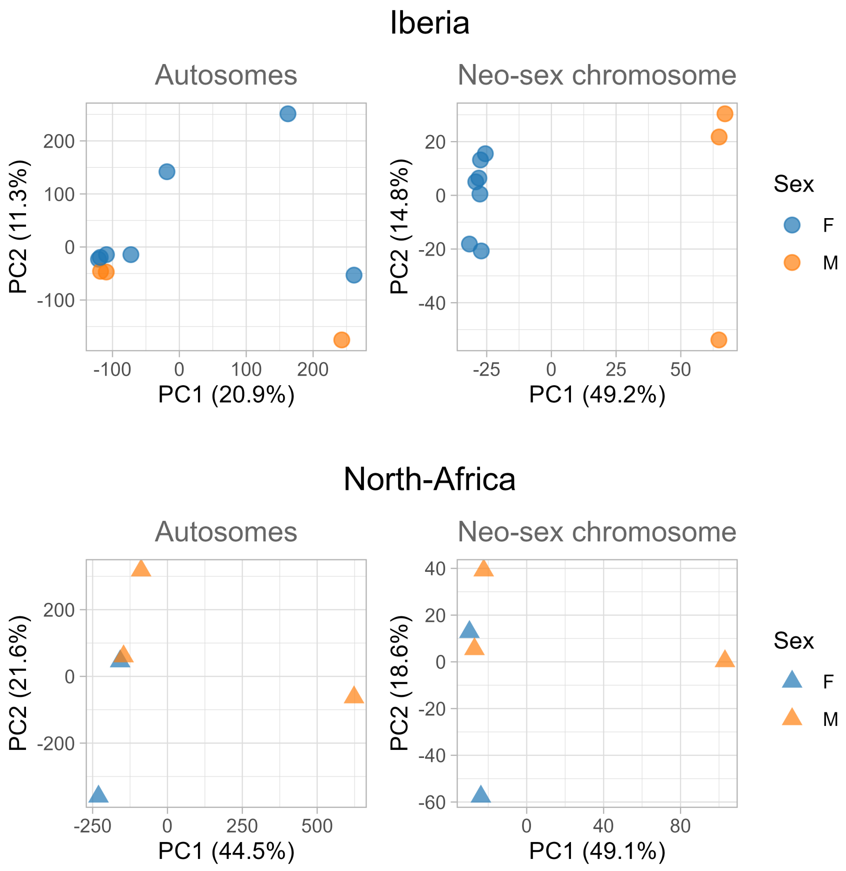


**Figure S2** – Detection of the neo-sex chromosome. PCA for the autosomes and chromosome 14, for samples from the Iberian peninsula and the Maghreb. Chromosome 14 clusters by sex for sample from the Iberian peninsula. By contrast, PCA on autosomes and chromosome 14 show similar patterns

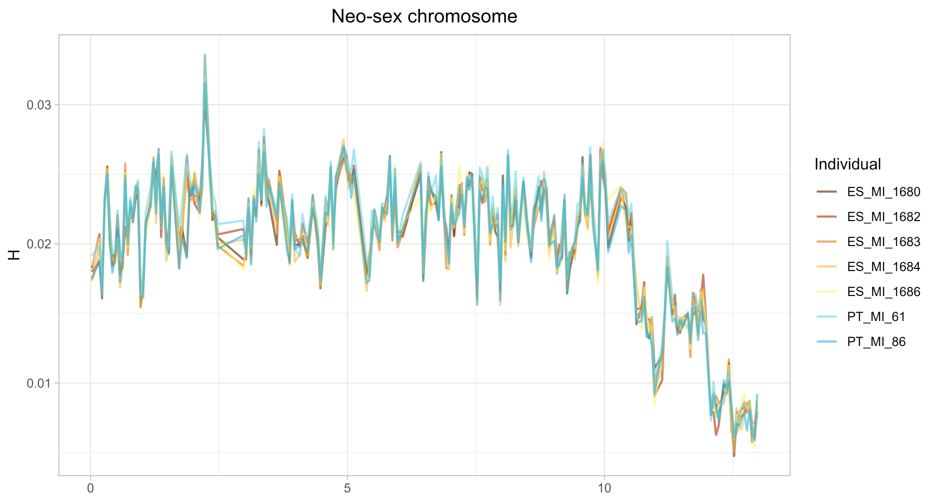


**Figure S3** – Windowed heterozygosity on the neo-sex chromosome of Iberian females

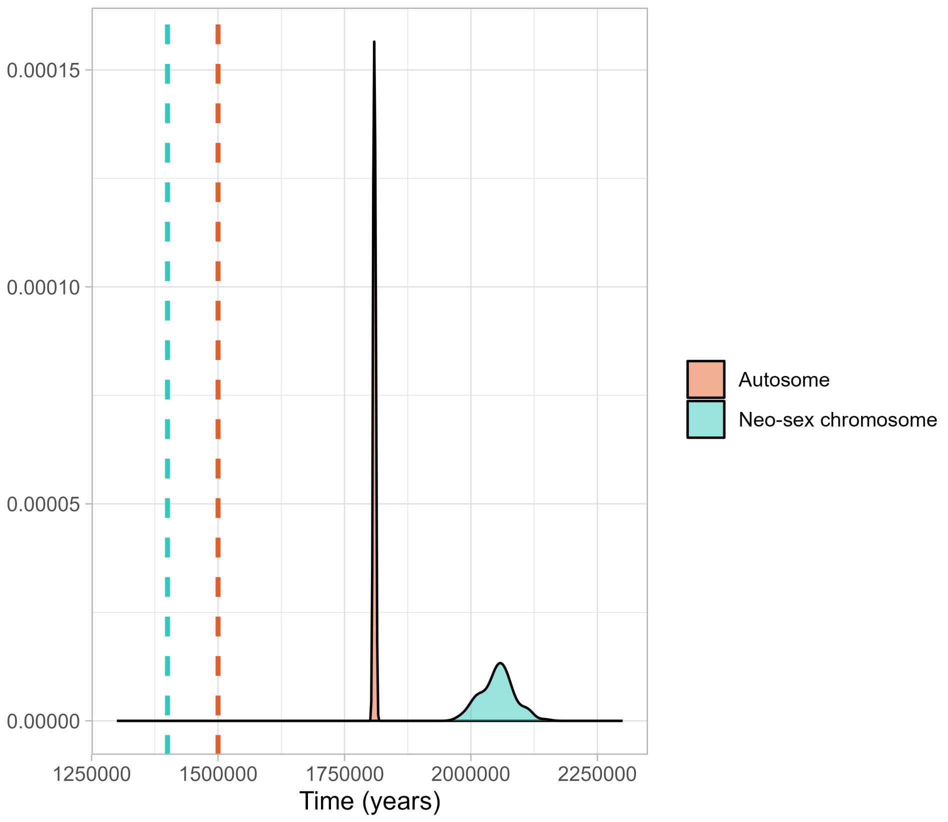


**Figure S4** – Density of maximum composite likelihood estimate for bootstrap replicates. Dashed lines are the maximum composite likelihood estimates of the original data

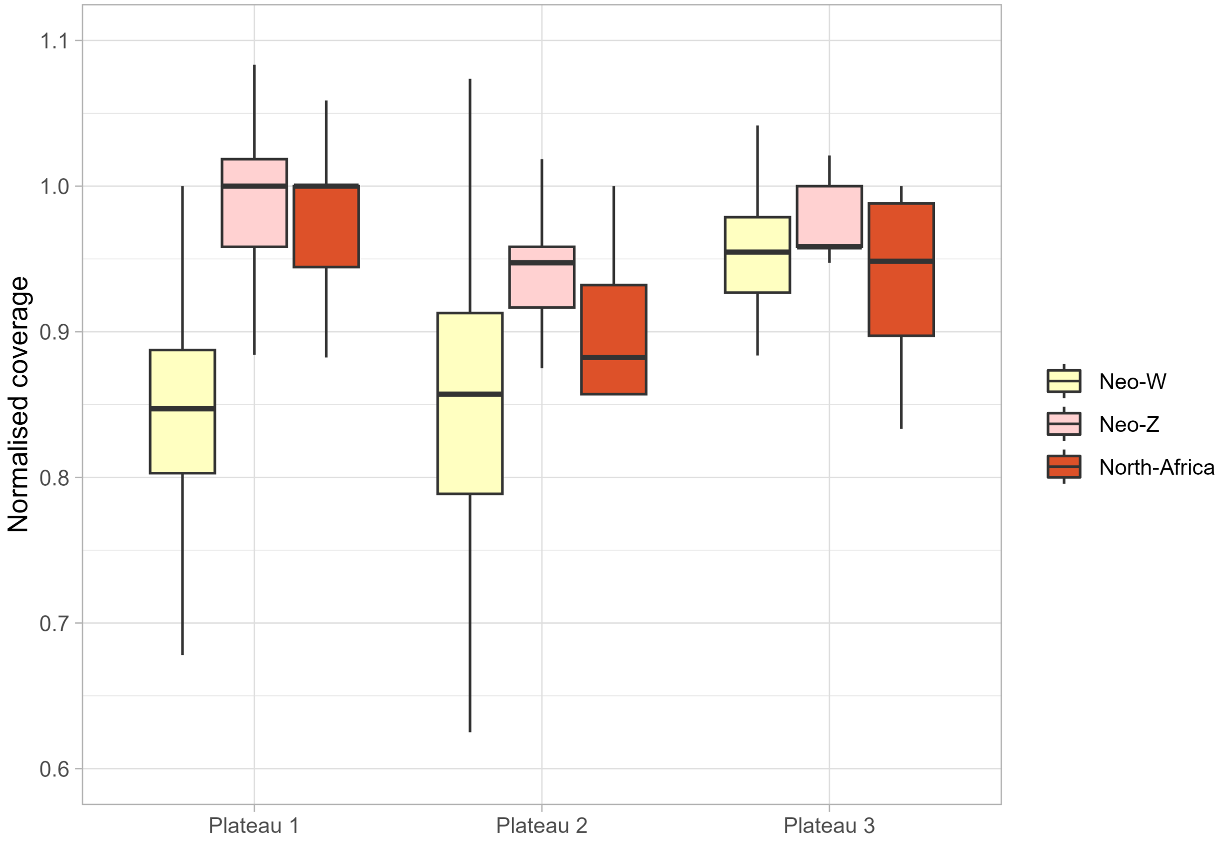


**Figure S5** – Relative coverage for the neo-Z and neo-W chromosomes, and their North-African homolog. Read depth was computed in windows of 100kb and normalised against the median autosomal coverage.

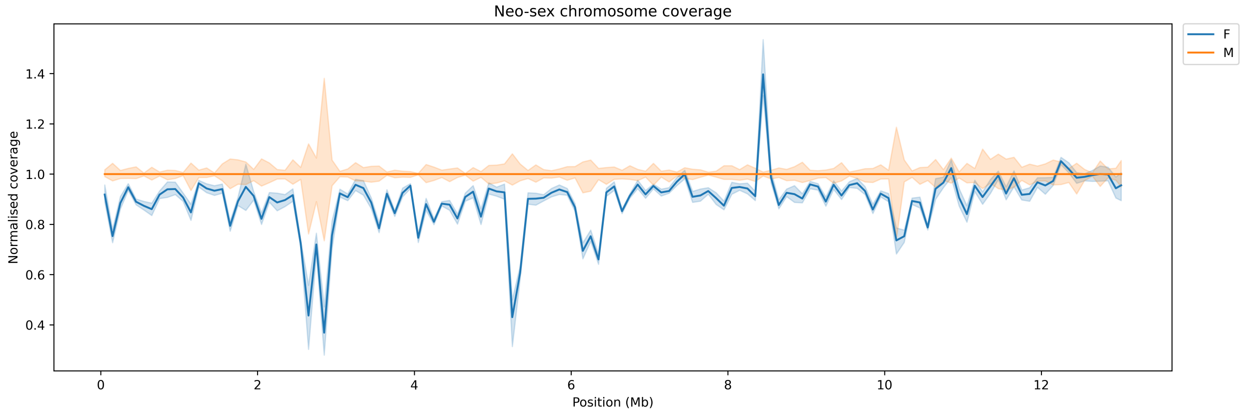


**Figure S6** – Relative coverage on the neo-sex chromosome for Iberian males and females. Read depth was computed in windows of 100kb and normalised against the median autosomal coverage.

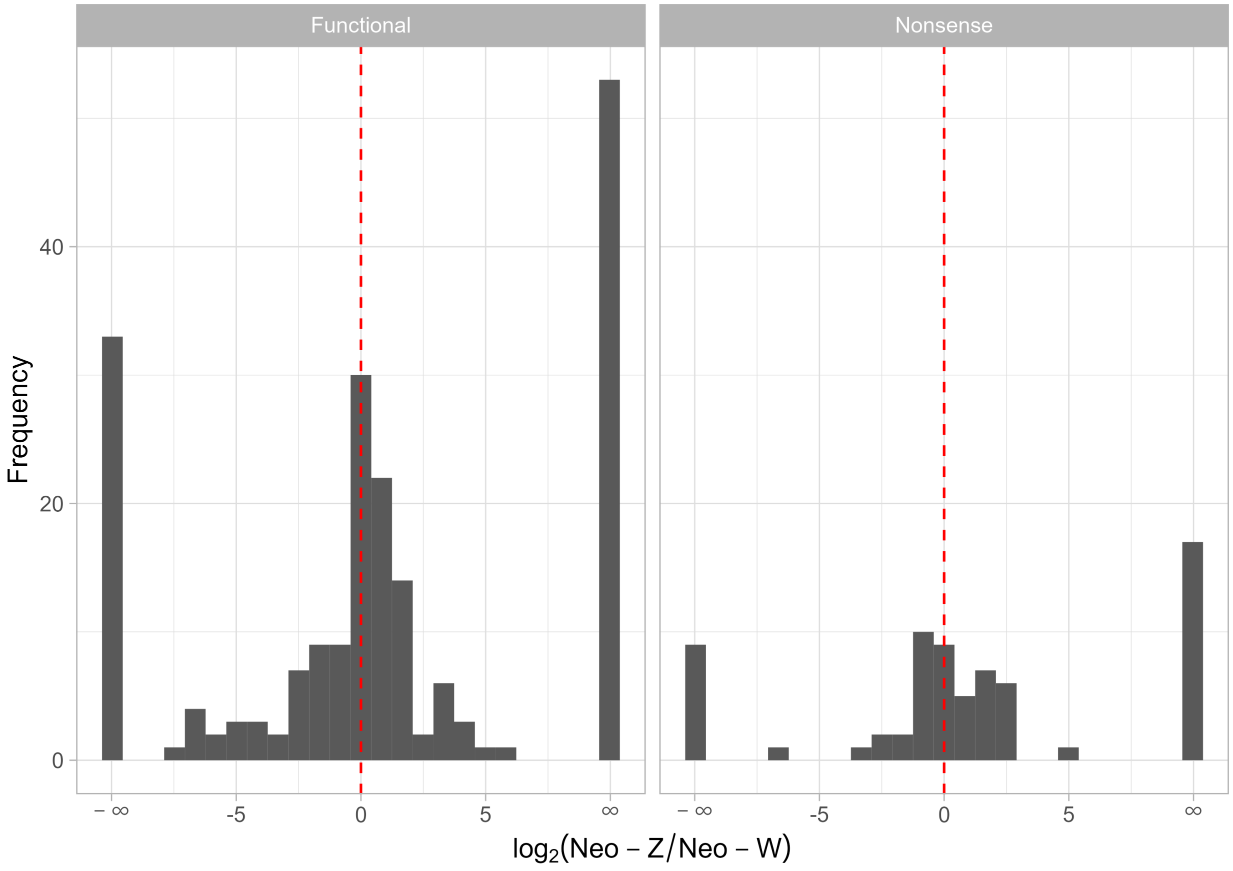


**Figure S7** – Relative expression of potentially functional and non-functional genes on the neo-Z and neo-W chromosomes.

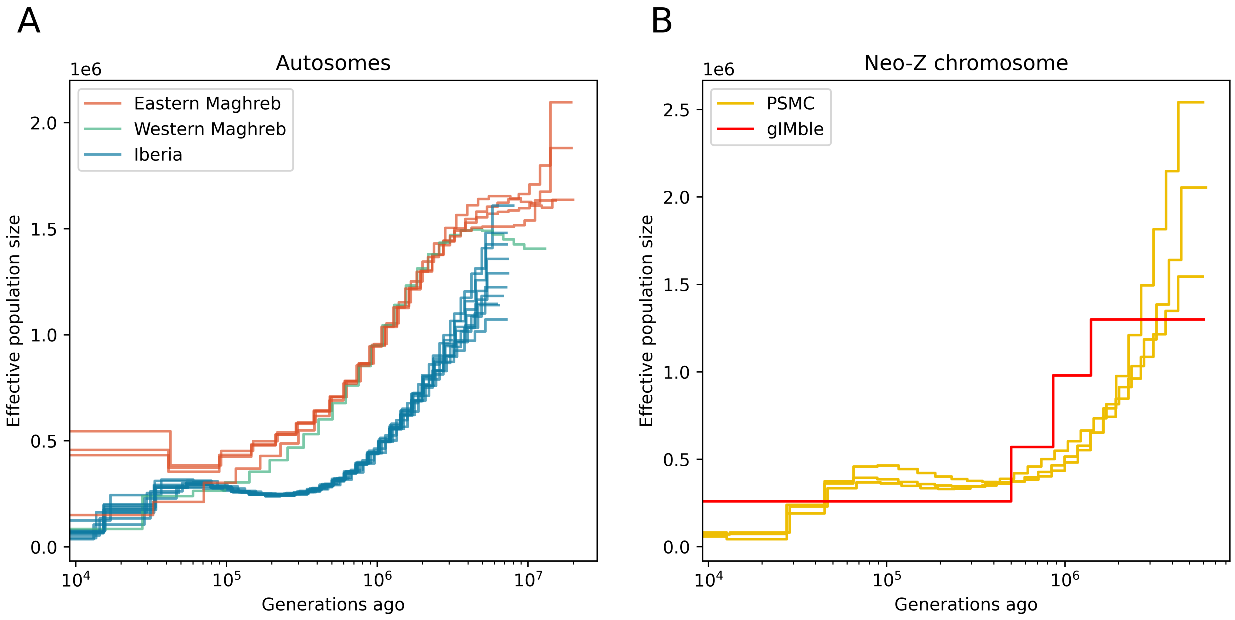


**Figure S8** – A) PSMC-inferred demographic history of samples from Iberia, Eastern and Western Maghreb. The three populations show similar patterns. B) Demographic history of the neo-Z chromosome, as inferred by PSMC for three Iberian males, and by gIMble using the ancestral population size and split times inferred for each plateaus. Reassuringly, the two inferred trajectories showed a close fit

**Table S1** – Sequenced individual metadata

| Sample | Preservation | Species | Sex | Country | Latitude | Longitude | Data |
|---|---|---|---|---|---|---|---|
| PT_MI_7 | Liquid nitrogen | *Melanargia ines* | M | Portugal | 37.67 | -7.738 | WGS |
| PT_MI_8 | Liquid nitrogen | *Melanargia ines* | M | Portugal | 37.67 | -7.738 | WGS, Pacbio |
| PT_MI_61 | Liquid nitrogen | *Melanargia ines* | F | Portugal | 37.211 | -7.438 | WGS |
| PT_MI_86 | Liquid nitrogen | *Melanargia ines* | F | Portugal | 37.411 | -7.802 | WGS, Hi-C |
| TN_MI_1619 | Ethanol | *Melanargia ines* | F | Tunisia | 35.876 | 8.547 | WGS |
| MA_MI_1620 | Ethanol | *Melanargia ines* | M | Morocco | 30.61 | -7.58 | WGS |
| DZ_MI_1624 | Ethanol | *Melanargia ines* | M | Algeria | 36.004 | 4.227 | WGS |
| ES_MI_1647 | Ethanol | *Melanargia ines* | M | Spain | 39.875 | -0.417 | WGS |
| ES_MI_1680 | Ethanol | *Melanargia ines* | F | Spain | 39.93 | -4.474 | WGS |
| DZ_MI_1681 | Ethanol | *Melanargia ines* | M | Algeria | 35.1 | 1.15 | WGS |
| ES_MI_1682 | Ethanol | *Melanargia ines* | F | Spain | 42.12639 | -1.39389 | WGS |
| ES_MI_1683 | Ethanol | *Melanargia ines* | F | Spain | 39.60081 | -7.18825 | WGS |
| ES_MI_1684 | Ethanol | *Melanargia ines* | F | Spain | 38.01498 | -1.37461 | WGS |
| DZ_MI_1685 | Ethanol | *Melanargia ines* | F | Algeria | 36.00083 | 4.297778 | WGS |
| ES_MI_1686 | Ethanol | *Melanargia ines* | F | Spain | 36.8426 | -5.35515 | WGS |

**Table S2** – Model fit. $\Delta \ln \mathrm{CL}$ measured relative to the best fitting model, $DIV$

| Model | $DIV$ | $IM_{\rightarrow \mathrm{EM}}$ | $IM_{\rightarrow \mathrm{IP}}$ | $MIG_{\rightarrow \mathrm{EM}}$ | $MIG_{\rightarrow \mathrm{IP}}$ |
|---|---|---|---|---|---|
| $\Delta \ln \mathrm{CL}$ | 0 | 0 | 0 | -5,607,055 | -4,004,366 |
